# Transcriptomic and metabolomic profiling reveals early responses of tomato plants elicited by compost-derived bacteria

**DOI:** 10.64898/2026.10.27.762366

**Authors:** Giulia Ghirardello, Fabrizio Araniti, Rosalba Cipriani, Werner Zanardi, Tiziano Bonato, Barbara Baldan, Sebastiano Nigris

## Abstract

Compost produced from biowaste has a defined microbiota, and some bacterial species may represent promising resources for low-input crop systems. However, their effects on plant physiology are poorly investigated and remain difficult to predict from standard bacterial traits alone. We isolated several bacterial strains (*Bacillus sonorensis, Glutamicibacter arilaitensis,* and *Kocuria rhizophila*) from biowaste compost and examined their early effects on tomato seedlings under controlled in vitro conditions. Responses were assessed by root phenotyping, root and shoot transcriptomics, and mass spectrometry-based profiling of primary metabolites in plant tissues, root exudates and bacterial culture filtrates. *B. sonorensis* induces the broadest response, reducing primary root length and lateral root number and density and triggering broad transcriptional changes related to metabolism, redox balance, hormone signalling and defence-associated pathways. *G. arilaitensis* elicits a more selective, tissue-dependent response, whereas *K. rhizophila* has limited phenotypic and transcriptional effects at this stage. All isolates reshape primary metabolism, with recurrent effects on amino acid-, carbohydrate- and lipid-related pathways, together with isolate-specific changes in root exudate composition. This study characterises the early molecular and metabolic responses of tomato to compost-derived bacteria and identifies common and strain-specific physiological signatures that provide a foundation for future studies under biotic and abiotic stress.

## Introduction

Achieving sustainable agriculture requires strategies that maintain crop productivity and soil health while reducing reliance on chemical fertilisers and pesticides. Soil microorganisms contribute to this goal by sustaining nutrient cycling, maintaining soil structure, suppressing disease and enhancing plant resilience to environmental perturbations [1]. Accordingly, microbiome-based approaches are increasingly considered promising tools for low-input and climate-constrained agriculture [2]. Compost from organic-waste (urban biowaste) recycling is widely used as an organic amendment that improves soil fertility and supports plant growth [3]. Similarly to compost from manure or from agricultural waste [4–6], compost from biowaste also harbours active microbial communities, including potential plant growth-promoting rhizobacteria (PGPR) [7–9]. PGPR can support plant performance through phosphate solubilisation, siderophore production, phytohormone modulation, pathogen antagonism and induced systemic resistance [10], but their effects on the plants are strongly strain-, host- and environment-dependent, and canonical PGP traits do not necessarily predict the outcome of a specific plant–bacterium interaction [11]. Compost-associated bacteria are mostly investigated at microbial-community levels [7], whereas the effects of individual compost-derived isolates remain less well resolved, and direct characterisation of these isolates is therefore needed to determine how their functional properties relate to plant responses [7,8]. Molecular techniques show considerable potential to characterize in depth the effects of direct inoculation of single strains on plant tissue. In fact, transcriptomics is increasingly combined with metabolomics to characterise plant responses to beneficial microorganisms [12–15]. Transcriptomic profiling identifies genes and molecular pathways associated with bacterial treatment, whereas metabolomics provides complementary information on the resulting biochemical and physiological state [16]. Primary metabolites are particularly informative because they connect growth with carbon and nitrogen allocation, redox balance and stress-associated metabolism [17]. Root exudates composition analysis provides an additional level of information because exudates mediate plant–microbe communication, influence microbial recruitment and reflect changes at the root chemical interface [18,19]. In addition, profiling bacterial culture filtrates can further characterise compounds released by individual isolates. Examining plant tissue transcriptional and metabolic response, root exudate and bacterial culture-filtrate profiles in parallel can therefore provide complementary perspectives on shared and isolate-specific plant responses and identify candidate processes for subsequent functional validation. In this study, we investigated three compost-derived bacterial isolates, *Bacillus sonorensis*, *Glutamicibacter arilaitensis* and *Kocuria rhizophila*, for their capacity to modulate early responses in tomato Micro-Tom seedlings . The isolates were previously characterised in vitro for phosphate solubilisation, siderophore production and indole-3-acetic acid biosynthesis as background for interpreting the plant-level responses that followed, and their effects on the plants were examined under controlled *in vitro* conditions. Root and shoot transcriptomic analyses were conducted in parallel with GC–MS-based profiling of primary metabolites in plant tissues, root exudates and bacterial culture filtrates, while root phenotyping provided a complementary assessment of early architectural responses. Here, we aimed to determine whether isolates displaying different *in vitro* Plant Growth Promotion-associated traits elicited common or isolate-specific plant responses, whether roots and shoots differed in their sensitivity to inoculation and whether the transcriptomic and metabolomic datasets revealed corresponding or distinct response patterns. This approach characterises the early molecular and metabolic states induced by individual compost-derived isolates and provides a basis for subsequent functional validation under soil-based and stress conditions.

## Results

### Bacteria inoculation causes different early root growth responses

Root phenotyping at 7 days post-inoculation does not reveal enhanced root growth under the tested conditions. *B. sonorensis* reduces primary root length, lateral root number, and lateral root density, whereas *G. arilaitensis* reduces lateral root number and density without significantly affecting primary root length. *K. rhizophila* does not significantly alter any of the measured root traits. Root diameter is not significantly affected by any treatment (Supplementary Fig. S1).

### *B. sonorensis* induces the broadest transcriptional response in shoots and roots

UpSet plots, which show the intersections among sets of differentially expressed genes (DEGs), reveal both treatment-specific and shared transcriptional responses (Fig. 1a, b). In shoots, 680 genes are uniquely regulated by *B. sonorensis* (494 upregulated and 186 downregulated), compared with 124 genes uniquely regulated by *G. arilaitensis* (121 upregulated and 3 downregulated) and 28 by *K. rhizophila* (13 upregulated and 15 downregulated). Fourteen genes, all upregulated, are shared among the three treatments (Fig. 1a). In roots, 766 genes are uniquely regulated by *B. sonorensis* (420 upregulated and 346 downregulated), 140 by *G. arilaitensis* (58 upregulated and 82 downregulated) and 78 by *K. rhizophila* (50 upregulated and 28 downregulated). A total of 94 genes are shared among the three root datasets, including 58 upregulated and 36 downregulated genes (Fig. 1b). The gene lists corresponding to each UpSet intersection, together with their KEGG annotations, are provided in Supplementary Tables S1–S14.

**Figure 1.**
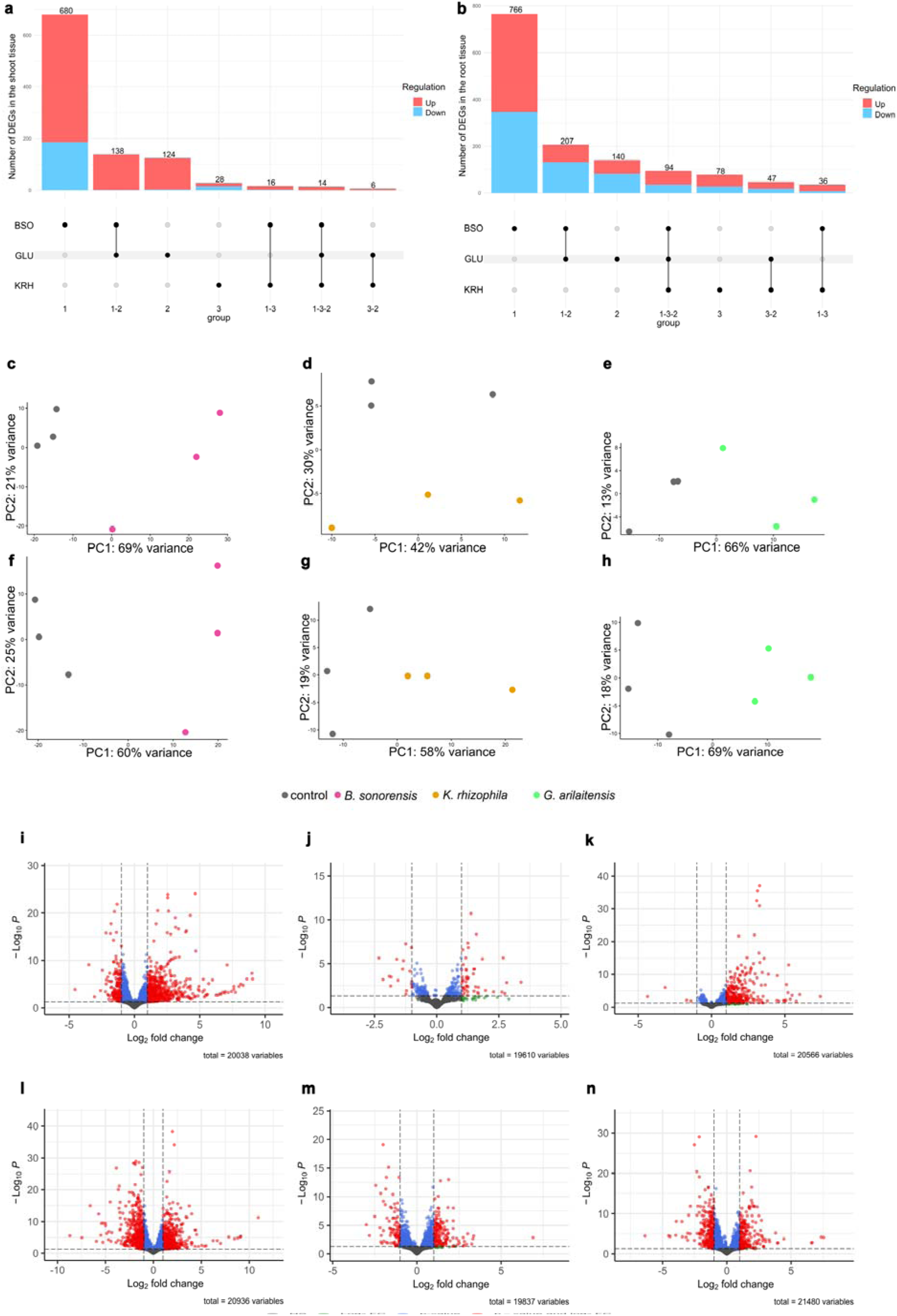
*Bacillus sonorensis* induces the broadest transcriptional response in tomato shoots and roots. (a,b) UpSet plots showing treatment-specific and shared differentially expressed genes (DEGs) in shoots (a) and roots (b) at 7 days post-inoculation. Red and blue fractions indicate upregulated and downregulated genes, respectively. Values above the bars indicate the sizes of the intersections defined by the connected dots below. BSO, *B. sonorensis*; KRH, *K. rhizophila*; GLU, *G. arilaitensis*. (c–h) Principal component analysis (PCA) of shoot (c–e) and root (f–h) transcriptomes from control samples and samples inoculated with BSO, KRH and GLU, respectively. n = 3 biological replicates per condition; each replicate comprised shoot or root tissue pooled from five seedlings. (i–n) Volcano plots showing differential gene expression in shoots (i–k) and roots (l–n) after treatment with BSO, KRH and GLU, respectively. The x-axis shows the log2 fold change, and the y-axis shows the −log10 adjusted p-value. Grey points meet neither threshold, green points meet only the |log2 fold change| ≥ 1 threshold, blue points meet only the adjusted p-value ≤ 0.05 threshold, and red points meet both thresholds. Significantly upregulated and downregulated genes are located on the right and left, respectively.

Principal component analysis (PCA) shows different degrees of separation between control and inoculated samples (Fig. 1c–h). In shoots, separation occurs mainly along PC1 for *B. sonorensis* and *G. arilaitensis*, with PC1 explaining more than 65% of the variance in both comparisons (Fig. 1c, e). Separation between control and *K. rhizophila*-treated shoots is less pronounced, with PC1 and PC2 explaining 42% and 30% of the variance, respectively (Fig. 1d). In roots, PC1 explains more than 60% of the variance for *B. sonorensis* and *G. arilaitensis* and 58% for *K. rhizophila* (Fig. 1f–h). PCA plots including all control and treated samples are shown in Supplementary Fig. S2.

Volcano plots show a predominance of upregulated DEGs in both shoots and roots (Fig. 1i–n). *K. rhizophila* elicits the weakest transcriptional response, particularly in shoots (Fig. 1j, m), whereas *B. sonorensis* produces the broadest distribution of significant DEGs in both tissues (Fig. 1i, l). Heatmaps summarising DEG expression patterns for each treatment and tissue are provided in Supplementary Fig. S3. The corresponding gene lists, ordered by heatmap rows and accompanied by KEGG annotations, are reported in Supplementary Tables S15–S20.

### *AP2/ERF* transcription factors recur across bacterial treatments and tissues

Differentially expressed transcription factor (TF) genes are identified in roots and shoots for each bacterial treatment (Supplementary Table S21). In roots, *B. sonorensis*, *K. rhizophila* and *G. arilaitensis* treatments are associated with 46, 23 and 34 upregulated TF genes and 73, 19 and 37 downregulated TF genes, respectively. In shoots, 73, 6 and 32 TF genes are upregulated in response to *B. sonorensis*, *K. rhizophila* and *G. arilaitensis*, respectively, whereas downregulated TF genes are detected only after *B. sonorensis* treatment (21 genes). Differentially expressed transcription factors common to all treatments in root and shoot are reported in Table 2.

**Table 1.** PGP traits of the bacteria studied. +/-: positive/negative response. P: phosphate; IAA: indole-3-acetic acid.

| Bacterial species | P solubilisation | Siderophore production | IAA biosynthesis |
| --- | --- | --- | --- |
| *B. sonorensis* | + | + | + |
| *K. rhizophila* | + | + | + |
| *G. arilaitensis* | - | + | + |

**Table 2.**
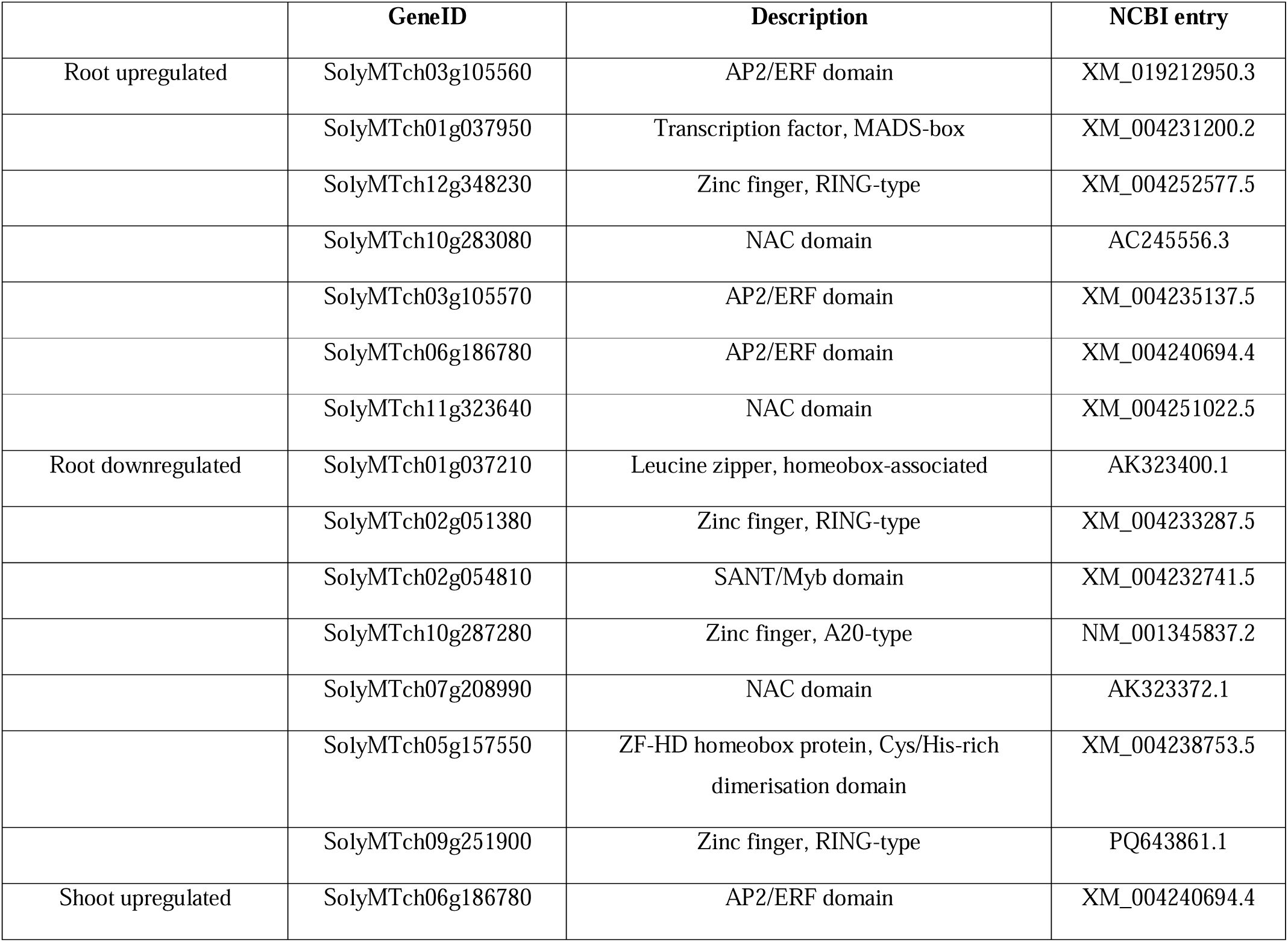
Differentially expressed genes encoding transcription factors found in common among the three plant treatments in the analysed tissues.

The AP2/ERF superfamily is the most represented class among upregulated root TFs, comprising 11 genes in *B. sonorensis*-treated roots, 7 in *K. rhizophila*-treated roots and 11 in *G. arilaitensis*-treated roots. Seven TF genes are upregulated in roots across all three treatments, including three AP2/ERF, two NAC, one MADS-box and one zinc finger RING-type gene. Seven TF genes are also commonly downregulated in roots and belong to homeobox-, zinc finger-, Myb- and NAC-related families. In shoots, a single AP2/ERF gene is upregulated by all three isolates. This gene is also commonly upregulated in roots, making it the only TF gene consistently upregulated across treatments and tissues.

### Descriptive KEGG mapping highlights a broad root response to *B. sonorensis*

Descriptive KEGG mapping, based on the net difference between up- and downregulated KEGG Orthology assignments, identifies tissue- and isolate-specific pathway patterns (Fig. 2a–f). The broad Metabolic pathways category is prominent in both tissues for all three isolates. It shows positive net DEG representation in *B. sonorensis*-treated shoots and roots (Fig. 2a, d), *G. arilaitensis*-treated shoots (Fig. 2c) and *K. rhizophila*-treated roots (Fig. 2e). Negative net representation is observed in *K. rhizophila*-treated shoots (Fig. 2b) and *G. arilaitensis*-treated roots (Fig. 2f).

**Figure 2.**
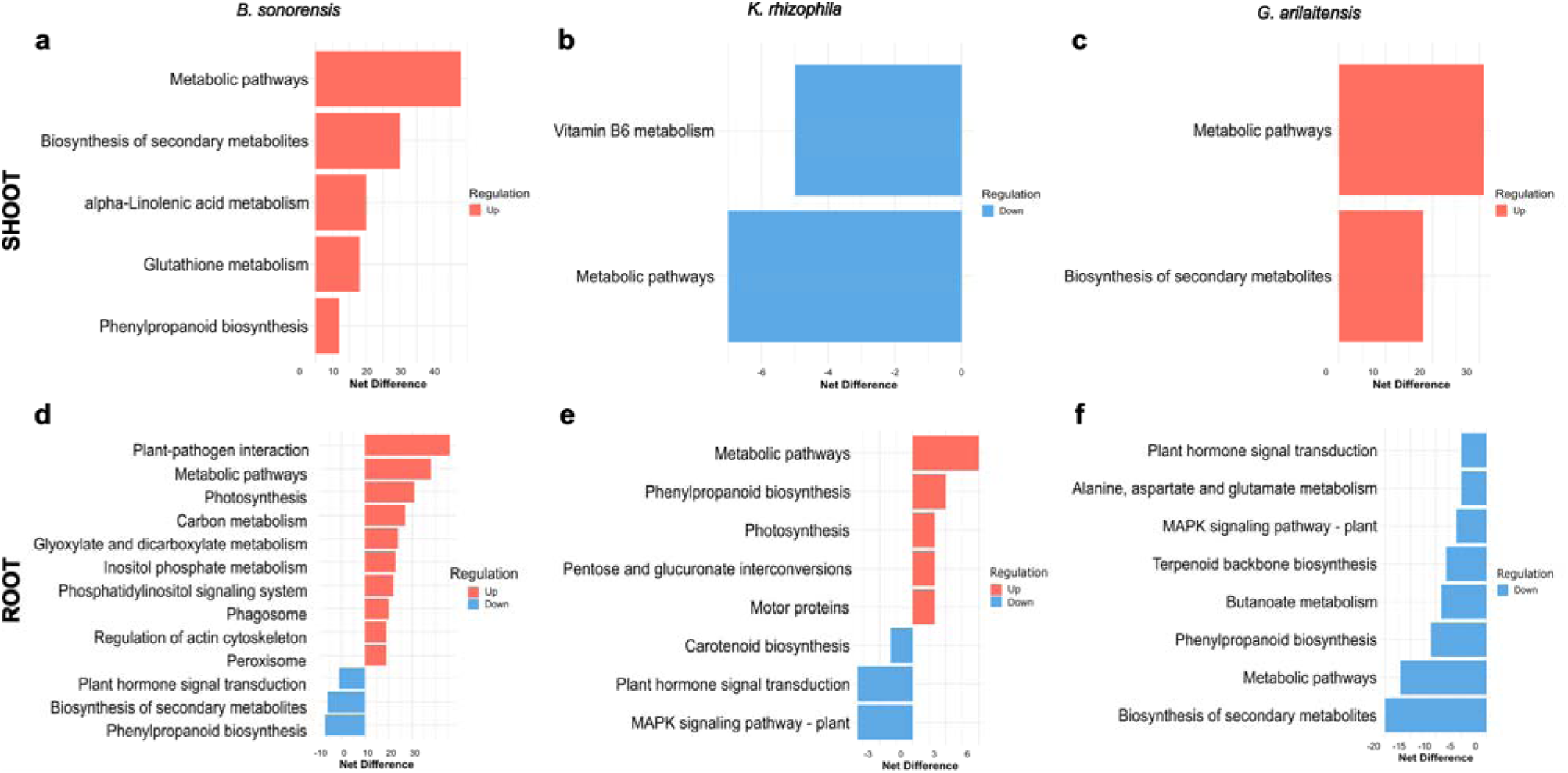
Descriptive KEGG mapping reveals tissue- and isolate-specific transcriptional patterns. KEGG categories are shown for shoots (a–c) and roots (d–f) at 7 days post-inoculation with *B. sonorensis*, *K. rhizophila* and *G. arilaitensis*, respectively. Net DEG representation was calculated as the number of KEGG Orthology assignments corresponding to upregulated genes minus the number corresponding to downregulated genes within each pathway. Red bars indicate positive net DEG representation, whereas blue bars indicate negative net DEG representation; these values do not indicate pathway activation or inhibition. Categories exceeding 1.5 times the standard deviation of the net-score distribution are shown. The corresponding thresholds are 6.7, 4.2 and 9.5 for panels a–c and 8.1, 2.1 and 4.6 for panels d–f. DEG, differentially expressed gene; KEGG, Kyoto Encyclopedia of Genes and Genomes.

In shoots, Biosynthesis of secondary metabolites shows positive net DEG representation after *B. sonorensis* and *G. arilaitensis* inoculation (Fig. 2a, c). *B. sonorensis* is also associated with positive representation of α-linolenic acid metabolism, Glutathione metabolism, and Phenylpropanoid metabolism, whereas Vitamin B6 metabolism shows negative representation only after *K. rhizophila* treatment (Fig. 2b).

Roots treated with *B. sonorensis* display the broadest set of prominent KEGG categories. Positive net DEG representation includes Plant–pathogen interaction, Photosynthesis, Carbon metabolism, Glyoxylate and dicarboxylate metabolism, Inositol phosphate metabolism, Phosphatidylinositol signalling system and Peroxisome, whereas Phenylpropanoid biosynthesis and Plant hormone signal transduction show negative representation (Fig. 2d). *K. rhizophila*-treated roots show positive representation of Phenylpropanoid biosynthesis, Photosynthesis, Pentose and glucuronate interconversions, and Motor proteins, together with negative representation of Carotenoid biosynthesis, Plant hormone signal transduction, and MAPK signalling pathway (Fig. 2e). *G. arilaitensis*-treated roots predominantly show negative net DEG representation, including Plant hormone signal transduction, Alanine, aspartate and glutamate metabolism, MAPK signalling pathway, Terpenoid backbone biosynthesis, Butanoate metabolism and Phenylpropanoid biosynthesis (Fig. 2f).

### Primary metabolite changes are widespread but differ between tissues and isolates

Heatmaps of significantly altered metabolites show that all three bacterial treatments affect the primary metabolite profiles of shoots and roots at 7 days post-inoculation. *B. sonorensis* alters 80 metabolites in shoots and 73 in roots, with most changes in roots corresponding to reduced abundance. *K. rhizophila* affects 66 metabolites in shoots and 62 in roots, whereas *G. arilaitensis* affects 78 metabolites in shoots and 76 in roots, with reductions predominating in shoots (Supplementary Fig. S4).

Orthogonal partial least squares–discriminant analysis (OPLS-DA), a supervised multivariate approach, shows separation between control and treated samples in all comparisons (Figs. 3a–c; 4a–c). The predictive component t1 explains more than 70% of the variation in comparisons involving *B. sonorensis* and *G. arilaitensis* and more than 50% in those involving *K. rhizophila*, in both shoots and roots. An overview including all treatments is provided in Supplementary Fig. S5. Metabolites contributing to group separation were selected using a variable importance in projection threshold of VIP > 1 (Supplementary Tables S22–S27) and are presented separately for shoots and roots in Figs. 3d–g and 4d–g.

**Figure 3.**
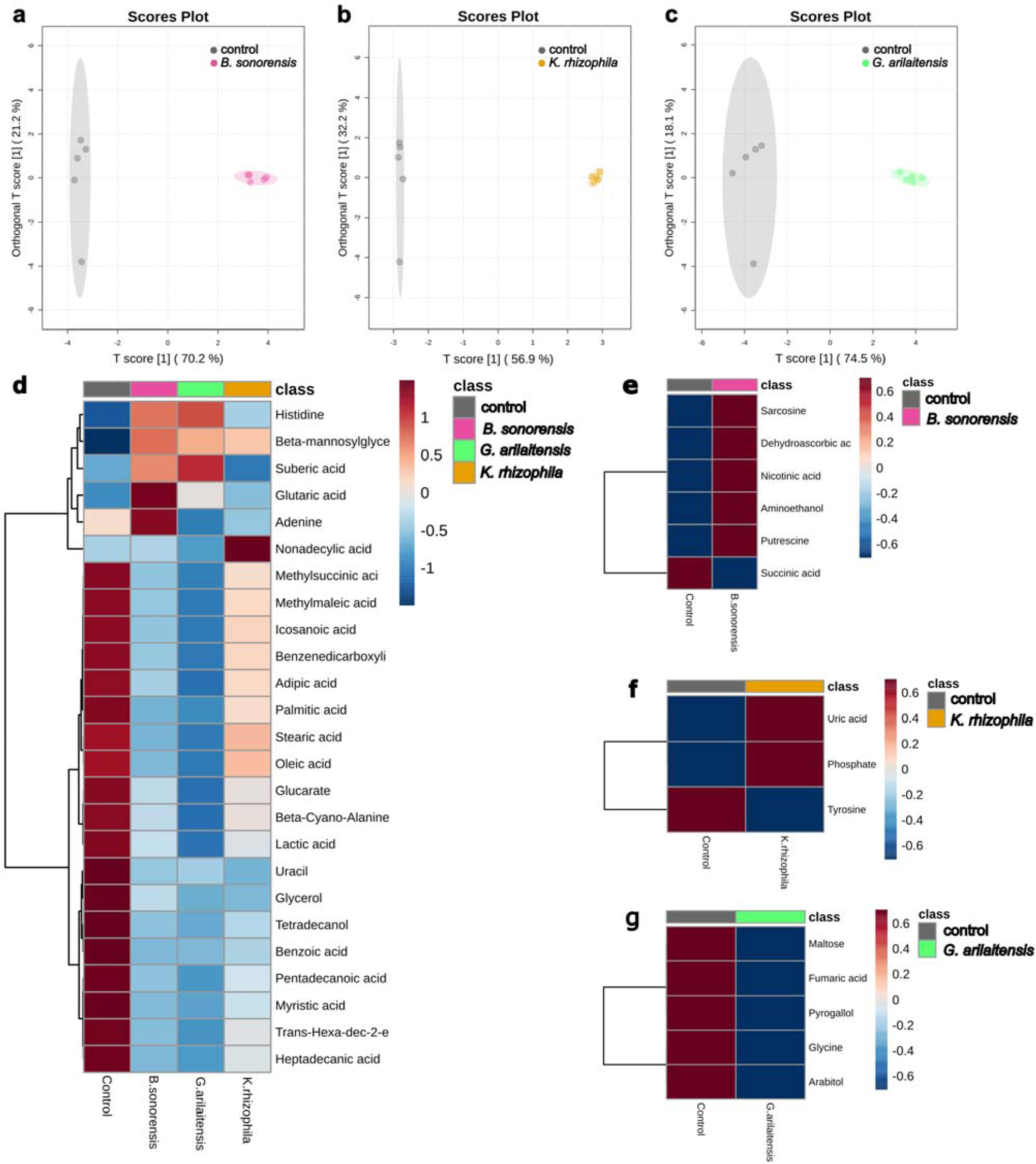
Bacterial inoculation generates distinct primary-metabolite profiles in tomato shoots. (a–c) Orthogonal partial least squares–discriminant analysis (OPLS-DA) score plots comparing control shoots with shoots treated with *B. sonorensis*, *K. rhizophila* and *G. arilaitensis*, respectively. The x- and y-axes show the predictive and orthogonal components, respectively, and shaded ellipses indicate the group confidence regions generated by the analysis. n = 5 biological replicates per condition; each replicate comprised shoot tissue pooled from five seedlings. (d–g) Heatmaps of metabolites with variable importance in projection (VIP) scores > 1. Panel d shows metabolites displaying a common response across the three bacterial treatments relative to the control. Panels e–g show treatment-specific metabolites for *B. sonorensis*, *K. rhizophila* and *G. arilaitensis*, respectively. Heatmap colours indicate relative metabolite abundance after data normalisation and scaling; columns represent condition means calculated from five biological replicates. Analyses were performed using MetaboAnalyst 6.0.

**Figure 4.**
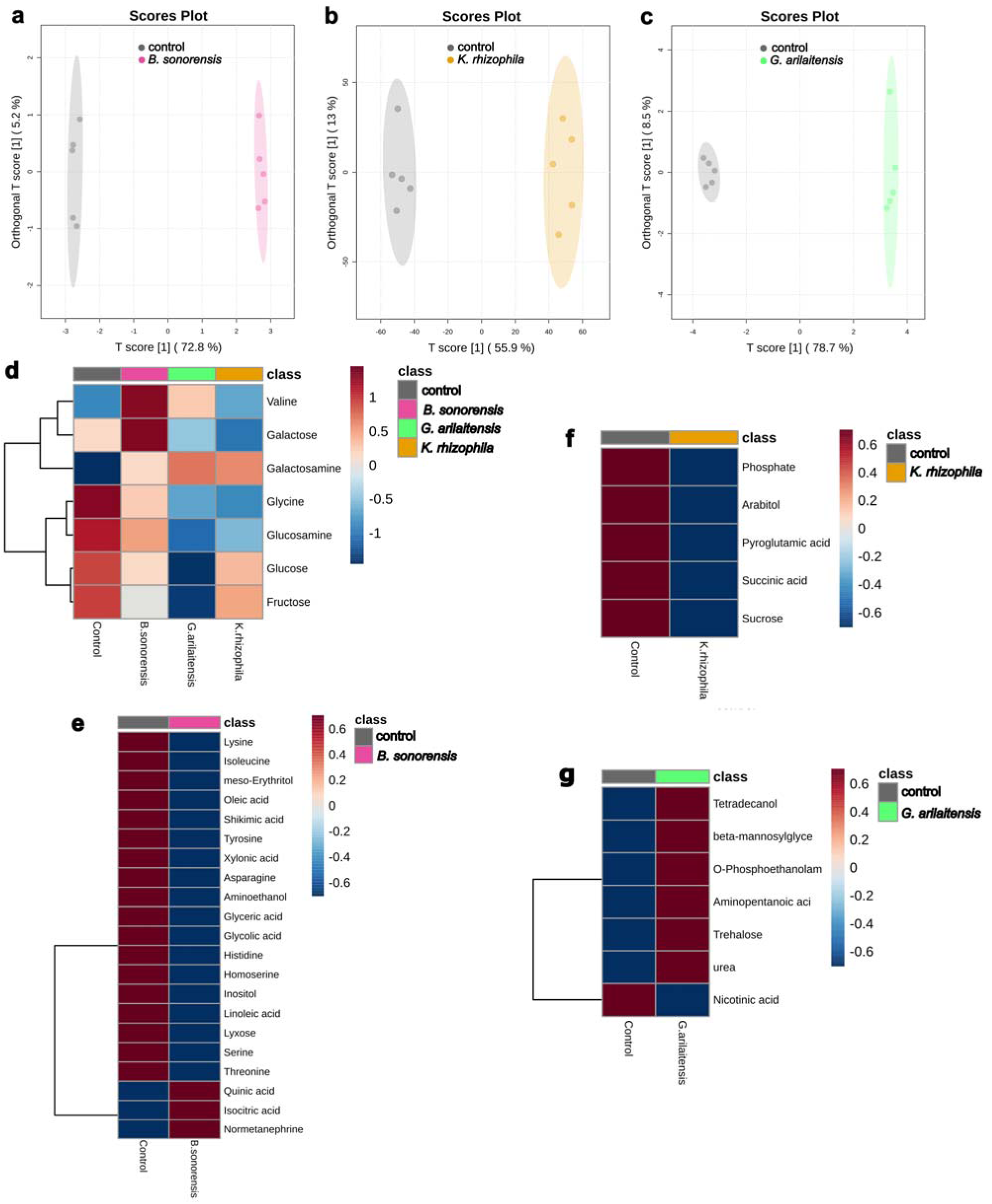
Bacterial inoculation generates distinct primary-metabolite profiles in tomato roots. (a–c) Orthogonal partial least squares–discriminant analysis (OPLS-DA) score plots comparing control roots with roots treated with *B. sonorensis*, *K. rhizophila* and *G. arilaitensis*, respectively. The x- and y-axes show the predictive and orthogonal components, respectively, and shaded ellipses indicate the group confidence regions generated by the analysis. n = 5 biological replicates per condition; each replicate comprised root tissue pooled from five seedlings. (d–g) Heatmaps of metabolites with variable importance in projection (VIP) scores > 1. Panel d shows metabolites displaying a common response across the three bacterial treatments relative to the control. Panels e–g show treatment-specific metabolites for *B. sonorensis*, *K. rhizophila* and *G. arilaitensis*, respectively. Heatmap colours indicate relative metabolite abundance after data normalisation and scaling; columns represent condition means calculated from five biological replicates. Analyses were performed using MetaboAnalyst 6.0.

In shoots, the three treatments share reductions in several fatty acids, organic acids, amino acids, sugars and nitrogen-containing compounds, including palmitic acid, stearic acid, lactic acid, histidine, β-cyanoalanine and adenine (Fig. 3d). Treatment-specific patterns are also observed. *B. sonorensis* increases aminoethanol, putrescine, sarcosine, dehydroascorbic acid and nicotinic acid, while reducing succinic acid (Fig. 3e). *K. rhizophila* increases uric acid and phosphate and reduces tyrosine (Fig. 3f). *G. arilaitensis* reduces maltose, arabitol, fumaric acid, pyrogallol and glycine (Fig. 3g).

In roots, all three treatments reduce several amino sugars, soluble sugars and amino acids, including galactosamine, glucosamine, glucose, fructose and glycine (Fig. 4d). Valine and galactose instead show isolate-dependent patterns. *B. sonorensis* produces the broadest treatment-specific response, predominantly reducing amino acids, organic acids, fatty acids, sugars and polyols, including serine, isocitric acid, linoleic acid and inositol (Fig. 4e). *K. rhizophila* reduces phosphate, arabitol, pyroglutamic acid, succinic acid and sucrose (Fig. 4f). By contrast, *G. arilaitensis* increases tetradecanol, β-mannosylglycerate, trehalose, aminopentanoic acid, O-phosphoethanolamine and urea, while reducing nicotinic acid (Fig. 4g). Metabolite abundance patterns across individual replicates are shown in Supplementary Figs. S6 and S7.

### Bacterial treatments converge on amino acid metabolism but differ in other pathways

Metabolic pathway analysis identifies recurrent effects on amino acid, carbohydrate and lipid metabolism (Table 3). In shoots, alanine, aspartate and glutamate metabolism, glycine, serine and threonine metabolism, and linoleic acid metabolism are significantly enriched across all three treatments, with impact scores of 0.78, 0.62 and 1.00, respectively. Starch and sucrose metabolism is significantly enriched only after *B. sonorensis* and *G. arilaitensis* treatment. In roots, alanine, aspartate and glutamate metabolism show the highest impact among the recurrently enriched pathways (0.85), followed by starch and sucrose metabolism (0.65) and glycine, serine and threonine metabolism (0.62). Linoleic acid metabolism has an impact score of 1.00 but is significantly enriched only in response to *B. sonorensis* and *G. arilaitensis*. Complete pathway lists, impact scores, and statistical values are provided in Supplementary Tables S28–S33.

**Table 3.** Metabolic pathways in shoots and roots of seedlings after the three bacterial treatments at 7 dpi, with an impact score > 0.5. Total cmpd: the total number of compounds in the pathway; Hits: number of compounds matched in the uploaded dataset; Raw p: the original p-value calculated from the enrichment analysis; FDR: false discovery rate; Impact: is the pathway impact value calculated from pathway topology analysis. Analyses performed with MetaboAnalyst 6.0. BSO, *B. sonorensis*; GLU, *G. arilaitensis*; KRH, *K. rhizophila*.

| Tissue | Pathway | Total cmpd | Hits | Impact | BSO |  | GLU |  | KRH |  |
| --- | --- | --- | --- | --- | --- | --- | --- | --- | --- | --- |
|  |  |  |  |  | Raw p | FDR | Raw p | FDR | Raw p | FDR |
| Shoot | Linoleic acid metabolism | 4 | 1 | 1.00 | 0.32106 | 0.00467 | 1.104E-07 | 3.21E-07 | 0.04027 | 0.04686 |
|  | Alanine, aspartate and glutamate metabolism | 22 | 8 | 0.78 | 0.037297 | 7.7E-04 | 1.107E-05 | 1.73E-05 | 5.809E-04 | 0.00115 |
|  | Starch and sucrose metabolism | 22 | 6 | 0.64 | 0.25963 | 0.33911 | 5.578E-04 | 6.491E-04 | - | - |
|  | Glycine, serine and threonine metabolism | 33 | 8 | 0.62 | 2.418E-04 | 1.55E-05 | 1.298E-08 | 6.39E-08 | 1.014E-06 | 5.010E-06 |
| Root | Linoleic acid metabolism | 4 | 1 | 1.00 | 0.007737 | 0.0082705 | 0.017117 | 0.017398 | - | - |
|  | Alanine, aspartate and glutamate metabolism | 22 | 8 | 0.85 | 8.033E-11 | 5.53E-10 | 1.992E-11 | 7.370E-11 | 1.605E-09 | 6.25E-09 |
|  | Starch and sucrose metabolism | 22 | 6 | 0.65 | 2.371E-05 | 3.27E-05 | 1.348E-07 | 2.59E-07 | 0.0015471 | 0.0019184 |
|  | Glycine, serine and threonine metabolism | 33 | 7 | 0.62 | 1.252E-09 | 3.91E-09 | 2.702E-09 | 5.58E-09 | 1.569E-07 | 3.04E-07 |

### All three isolates remodel root exudation through distinct chemical signatures

Root exudate profiling reveals distinct responses to the three bacterial treatments. Relative to the control, *B. sonorensis*, *K. rhizophila* and *G. arilaitensis* alter the abundance of 69, 74 and 56 metabolites, respectively, with increases predominating in all treatments (Supplementary Fig. S8).

OPLS-DA shows separation between control and treated exudate samples in all comparisons (Fig. 5a–c). The predictive component t1 explains approximately 75% of the variation for *B. sonorensis*, 85% for *K. rhizophila* and 72% for *G. arilaitensis*. An overview including all treatments is provided in Supplementary Fig. S9. Metabolites contributing to group separation were selected using a VIP > 1 threshold (Supplementary Tables S34–S36) and are shown in Fig. 5d–g.

**Figure 5.**
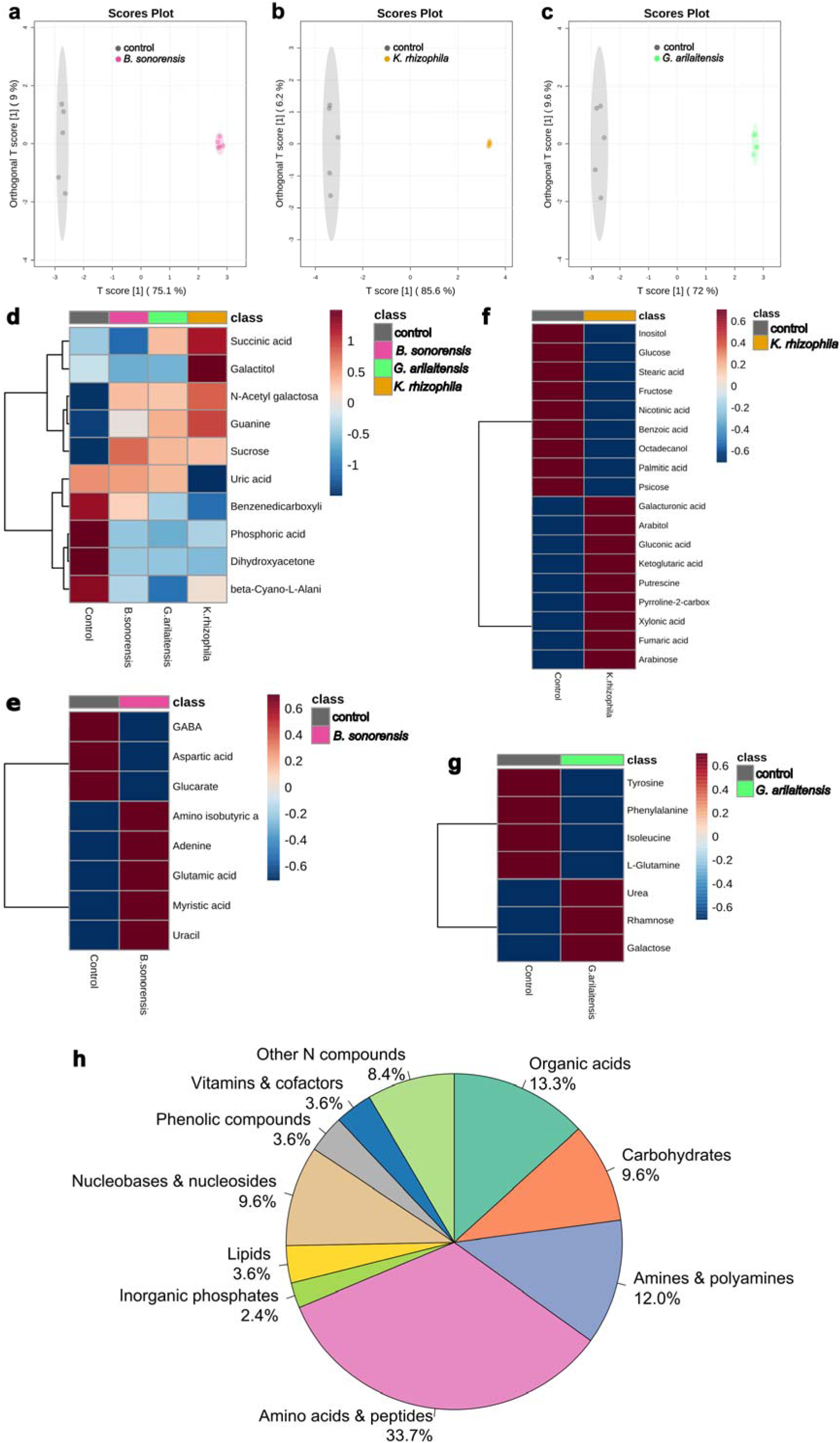
Bacterial treatments induce distinct changes in tomato root exudate profiles, while culture filtrates contain a broad range of metabolite classes. (a–c) Orthogonal partial least squares–discriminant analysis (OPLS-DA) score plots comparing root exudates from control seedlings with those from seedlings treated with *Bacillus sonorensis*, *Kocuria rhizophila* and *Glutamicibacter arilaitensis*, respectively. The x- and y-axes show the predictive and orthogonal components, respectively, and shaded ellipses indicate the group confidence regions generated by the analysis. n = 5 biological replicates per condition, each replicate comprised root exudates collected from five seedlings. (d–g) Heatmaps of root-exudate metabolites with variable importance in projection (VIP) scores > 1. Panel d shows metabolites displaying a common response across the three bacterial treatments relative to the control. Panels e–g show treatment-specific metabolites for *B. sonorensis*, *K. rhizophila* and *G. arilaitensis*, respectively. Heatmap colours indicate relative metabolite abundance after data normalisation and scaling; columns represent condition means calculated from five biological replicates. (h) Chemical-class distribution of the 83 annotated compounds detected across the culture filtrates of the three bacterial isolates. Percentages indicate the proportion of annotated compounds assigned to each chemical class. Analyses shown in panels a–g were performed using MetaboAnalyst 6.0.

Several metabolites show common responses across treatments. Benzenedicarboxylic acid, phosphoric acid, β-cyano-L-alanine and dihydroxyacetone are reduced in all inoculated samples, while guanine, N-acetylgalactosamine and sucrose are most abundant in the control (Fig. 5d). Other compounds display isolate-dependent patterns. Galactitol increases after *K. rhizophila* and *G. arilaitensis* treatment but decreases after *B. sonorensis* treatment. Succinic acid is reduced by *B. sonorensis* and increased by the other two isolates, whereas uric acid reaches its highest abundance in *K. rhizophila*-treated samples.

Treatment-specific profiles are also detected. *B. sonorensis* reduces GABA, aspartic acid and glucarate while increasing adenine, uracil, glutamic acid and myristic acid (Fig. 5e). *K. rhizophila* reduces several soluble sugars and lipid-related compounds and increases putrescine and selected organic and sugar acids (Fig. 5f). *G. arilaitensis* reduces several amino acids while increasing urea, rhamnose and galactose (Fig. 5g). Metabolite abundance patterns across individual replicates are shown in Supplementary Fig. S10.

### Bacterial culture filtrates reveal isolate-specific metabolic fingerprints

Metabolite profiling of cell-free culture filtrates reveals distinct chemical profiles for *B. sonorensis*, *K. rhizophila* and *G. arilaitensis*. Principal component analysis (PCA) clearly separates the three bacterial cultures, with PC1 and PC2 explaining 79.8% and 15.7% of the total variance, respectively (Fig. S11). The corresponding loadings plot identifies isocitric acid, xylitol, pyrrole-2-carboxylic acid, and threonic acid among the metabolites contributing most strongly to sample separation (Fig. S11).

Hierarchical clustering of metabolites showing significant differences by one-way ANOVA further highlights isolate-specific profiles (Fig. S11). *K. rhizophila* filtrates show comparatively higher levels of a metabolite cluster including ornithine, mannose, sorbitol and benzoic acid derivatives, whereas several amino acids and related compounds, including tyrosine, phenylalanine, methionine and tryptophan, are more abundant in *B. sonorensis* and *G. arilaitensis* filtrates. Complete metabolomic and statistical results for the bacterial cultures are provided in Supplementary File 1.

Among the 83 annotated compounds, amino acids and peptides represent the largest chemical class (33.7%), followed by organic acids (13.3%), amines and polyamines (12.0%), carbohydrates (9.6%) and nucleobases and nucleosides (9.6%). Other nitrogen-containing compounds account for 8.4%, while lipids, phenolic compounds, and vitamins and cofactors each represent 3.6%; inorganic phosphates account for the remaining 2.4% (Fig. 5h). The complete list of compounds is provided in Supplementary File 1

## Discussion

Inoculation with three compost-derived isolates, *B. sonorensis*, *G. arilaitensis* and *K. rhizophila*, triggers distinct early transcriptomic and metabolomic reprogramming in tomato roots and shoots. Although *B. sonorensis* and *K. rhizophila* share positive phosphate solubilisation, siderophore production and IAA biosynthesis *in vitro*, they induce markedly different molecular and metabolic profiles *in planta*. Conversely, *G. arilaitensis* shows the highest IAA production but does not elicit the broadest plant response, and root phenotyping at 7 dpi does not reveal enhanced root growth under the tested conditions. This dissociation between in vitro PGP-associated traits and the magnitude or direction of the plant response indicates that such traits offer, at best, a preliminary indication of functional potential rather than a predictor of the early molecular state induced in the plant [20]. Compost-derived isolates therefore remain a relevant source of candidate microorganisms for sustainable crop management, but their effects on plants need to be resolved directly rather than inferred from standard in vitro traits or community-level studies [7,8].

Across the three treatments, primary metabolism emerges as a major target of the early interaction, whereas defence- and signalling-related pathways are prominent only in specific isolate–tissue combinations. Metabolic pathway analysis recurrently highlights alanine, aspartate and glutamate metabolism; glycine, serine and threonine metabolism; starch and sucrose metabolism; and linoleic acid metabolism. Together, these pathways indicate coordinated adjustments in carbon–nitrogen balance, photorespiratory and one-carbon metabolism, carbohydrate allocation, fatty-acid-dependent membrane remodelling and oxylipin- and jasmonate-related signalling [21–24]. Their modulation is particularly strain-specific in roots, where decreases in sugars, amino acids, fatty acids and organic acids are more frequent than in shoots. Thus, these data identify common metabolic nodes affected by bacterial inoculation, and are consistent with the fact that direction and magnitude of metabolic changes reflect the tissue under examination and the physiological state associated [17].

Transcription factor patterns provide a complementary view of these responses. SolyMTch06g186780, annotated as an ERF025-like AP2/ERF and corresponding to SlERF5 in our analysis, is the only TF gene upregulated by all three isolates in both roots and shoots, suggesting a shared ethylene-related regulatory component. In roots, the common upregulated signature also includes the AP2/ERF gene SlERF.B5 and the NAC-domain gene SlNAC90. SlERF5 has been associated with tolerance to drought and salinity and resistance to *Botrytis cinerea*, whereas SlERF.B5 contributes to ascorbic acid biosynthesis, ROS scavenging and tolerance to oxidative and salt stress [25–27]. SlNAC90 participates in a jasmonate-regulatory module involved in the tomato response to low temperature [28]. Together, these patterns suggest that the treatments converge on regulatory components related to hormone signalling, redox control and stress-associated responses. The induction of SlAGL23 is more difficult to interpret because this MADS-box gene has mainly been associated with female gametophyte and embryo development and has not, to our knowledge, been characterised in roots or plant–microbe interactions [29,30]. Among the commonly downregulated root TFs, SlJA2L is a NAC-domain transcription factor involved in JA- and coronatine-dependent stomatal reopening during tomato–pathogen interactions through the regulation of salicylic acid metabolism [31]. Its downregulation suggests that the response to these isolates does not simply mirror this pathogen-associated JA response. Overall, the shared TF patterns identify candidate regulatory markers for future functional studies testing the transcriptional response under biotic and abiotic stresses

Within this common framework, *B. sonorensis* elicits the broadest transcriptional and metabolic response in both tissues and induces the highest number of differentially expressed TF genes. The root response is particularly broad at the KEGG level. Positive net DEG representation in Plant–pathogen interaction occurs together with Carbon metabolism, Peroxisome, Inositol phosphate metabolism and Phosphatidylinositol signalling system. This pattern indicates that the response is not limited to primary metabolic changes but also includes genes associated with microbial perception, redox regulation and membrane-related signalling. The Photosynthesis category detected in roots is interpreted as a KEGG annotation reflecting plastid- and carbon-associated genes, rather than as evidence of photosynthetic activation in root tissue.

Genes assigned to Plant–pathogen interaction include components commonly associated with microbial perception and immune signalling, including elements of PTI/ETI-related pathways. Under the sterile and non-stressed conditions used here, this pattern is consistent with defence-related signalling during early root–bacterium recognition, although it does not demonstrate activation of a pathogen-like response. The coexistence of defence-associated signatures, hormone-related categories and broad metabolic changes is consistent with the extensive crosstalk among salicidic acid, jasmonic acid, Ethylene, auxin, gibberellin and brassinosteroid pathways that coordinates plant growth and responses to external stimuli [32–34].

The metabolome of *B. sonorensis*-treated roots shows the strongest depletion across amino acids, organic acids, fatty acids and carbohydrate-related compounds. Reductions in serine, isocitric acid, linoleic acid and inositol are consistent with changes involving amino-acid metabolism, TCA-linked organic acids, lipid remodelling and phosphoinositide-associated signalling. Together with the representation of glutathione-, lipid- and defence-related categories, these changes support an early reorientation of plant physiology towards redox regulation, membrane-associated processes and defence-related functions. The broad transcriptomic and metabolomic reprogramming induced by this isolate, spanning defense-related signalling, redox regulation and carbon–nitrogen metabolism, indicates an early molecular state distinct from what has been reported for other *B. sonorensis* isolates, which have instead been characterized through root-growth and biocontrol phenotypes under salinity and Fusarium challenge [35]. This divergence underlines that the molecular response is strain- and condition-specific, and provides a basis for testing whether this early signature predicts a functional outcome under subsequent stress or pathogen challenge*. G. arilaitensis* defines a more selective and organ-specific configuration. In shoots, the response indicates selective modulation of central metabolism, with changes in compounds related to carbon and amino-acid metabolism, including maltose, fumaric acid and glycine. In roots, the prevailing transcriptomic pattern shows negative net representation of hormone-, MAPK- and phenylpropanoid-related categories, while the metabolomic data indicate adjustments in amino-acid, nitrogen and carbohydrate metabolism. Treatment-specific increases in trehalose, urea and O-phosphoethanolamine further suggest changes in osmotic balance, nitrogen status and membrane-associated metabolism. Unlike *B. sonorensis*, this isolate does not show broad positive net representation of defence-related categories but instead induces a more selective reorganisation of signalling and metabolism. Such a configuration may be relevant under stress conditions such as salinity, consistent with reports that show *Glutamicibacter* spp. improve salt tolerance in rice and halophytic species [36,37].

*K. rhizophila* elicits the least extensive overall response at 7 dpi under the present *in vitro* conditions, although changes remain detectable in both transcriptomic and metabolomic datasets. It produces the lowest number of DEGs, while roots show changes in categories related to phenylpropanoid metabolism, hormone signalling, MAPK signalling and carbohydrate-related processes. The metabolomic response is similarly selective: in shoots, *K. rhizophila* increases uric acid and phosphate while reducing tyrosine, whereas in roots it reduces sucrose, succinic acid and phosphate. Overall, this profile indicates limited broad-scale reorganisation but detectable adjustments in signalling and carbon- and phosphate-associated processes. in addition to what we observed, it is also known that *K. rhizophila* increase tomato biomass and induce coordinated transcriptomic, proteomic and metabolic changes after longer-term cultivation [38]. Other *K. rhizophila* strains have also improved salt and water-related stress tolerance in maize and pea [39]. These comparisons emphasise the influence of the isolate, host, experimental system and sampling time.

Root exudate profiling extends the analysis from internal plant metabolism to the plant–bacterium interface. All three treatments modify exudate composition, generating isolate-specific patterns across sugars, amino-acid-related compounds, organic acids, lipids and nitrogenous metabolites. These changes may alter the availability of substrates and signals at the root surface and thereby influence bacterial colonisation or rhizosphere niche formation [19]. In *B. sonorensis*-treated seedlings, reductions in succinic acid, GABA and aspartic acid, together with increases in glutamic acid and myristic acid, indicate a marked shift in carbon-, nitrogen- and lipid-related compounds. Organic acids in tomato root exudates can influence *Bacillus* colonisation [40]. *K. rhizophila* treatment reduces soluble sugars and inositol while increasing organic and sugar acids, indicating altered carbon availability and organic-acid-mediated interactions. Plant myo-inositol transport can influence bacterial colonisation phenotypes, although the significance of its reduction in the present system remains uncertain [41]. *G. arilaitensis* reduces several amino acids and increases urea, rhamnose and galactose, indicating a shift towards nitrogen- and sugar-containing compounds at the root surface. These isolate-specific profiles describe changes in the chemical interface, but their ecological consequences cannot be determined under the present *in vitro* conditions.

Metabolomic analysis of bacterial culture filtrates reveals distinct metabolic fingerprints among the three isolates. These differences may help contextualise the distinct plant transcriptomic, tissue-metabolite and exudate profiles. The detection of IAA is consistent with the auxin-biosynthesis trait identified in the PGP assays, whereas spermidine and pipecolic acid are compounds potentially related to plant growth and defense responses. Bacterially produced spermidine has been associated with plant growth promotion in another *Bacillus*–plant system [42], while pipecolic acid participates in defence amplification and systemic acquired resistance in plants [43]. However, their detection in bacterial filtrates does not establish equivalent functions in the present interaction. The filtrates also contain hydroxybenzoic acids and several organic acids, including isocitric, lactic, malic and saccharic acids, which contribute to the distinct chemical environment generated by each isolate and may be relevant to nutrient mobilisation and rhizosphere chemistry. Because several compounds occur in bacterial filtrates, plant tissues and root exudates, their source cannot be assigned from the present data. Quantitative approaches are therefore needed to distinguish plant-derived from bacterial-derived metabolites and to establish causal relationships between specific bacterial compounds and plant responses. Overall, culture-filtrate profiling provides useful chemical context for interpreting the physiological responses induced by the three isolates.

This study shows that three compost-derived bacterial isolates establish distinct early interaction profiles in tomato seedlings that cannot be fully predicted by canonical *in vitro* PGP traits. Parallel transcriptomic and metabolomic analyses identify *B. sonorensis* as the isolate inducing the broadest response in both roots and shoots, whereas *G. arilaitensis* produces a more selective and tissue-dependent configuration and *K. rhizophila* elicits the least extensive, but still detectable, response. All three isolates affect primary metabolism and modify root exudate composition, while their culture filtrates display distinct chemical profiles.

The use of young seedlings under controlled *in vitro* conditions and the analysis of a single early time point limit the association of these molecular and metabolic signatures with stable growth-promoting or stress-protective effects. Nevertheless, the identified pathways, transcription factors and metabolites provide a basis for subsequent functional validation under soil-based conditions and defined biotic or abiotic stresses.

## Methods

### Plant material and growth conditions

Seeds of *Solanum lycopersicum* cv. Micro-Tom from USDA National Plant Germplasm System GRIN-global (Accession #PI 539919) were surface-sterilised in 5% (v/v) sodium hypochlorite for 20 min, rinsed four times with autoclaved deionised water, and incubated overnight in autoclaved tap water with gentle agitation to synchronise germination. Seeds were then sown on solid ½ MS (half-strength Murashige and Skoog basic medium) adjusted to pH 5.5 in square plastic plates, sealed, and grown vertically in a controlled-environment chamber (25 °C, 68% relative humidity, 300 µmol m□² s□¹ photosynthetic photon flux density, 16/8 h light/dark).

### Bacterial strains and PGP trait assays

Bacterial strains were previously isolated from compost produced from biowaste by S.E.S.A. SpA. The strains have been identified as *Bacillus sonorensis*, *Kocuria rhizophila*, and *Glutamicibacter arilaitensis* based on whole-genome assembly data. Plant growth-promoting traits (IAA production, phosphate solubilization, and siderophore production were assessed and reported here (Table 1) from Cipriani et al. (unpublished data). *Bacillus sonorensis* and *Kocuria rhizophila* test positive for all three traits, whereas *Glutamicibacter arilaitensis* produces siderophores and IAA but does not solubilise phosphate under the tested conditions. IAA concentrations differ among the isolates, reaching 1.615 µg mL□¹ in *B. sonorensis*, 5.985 µg mL□¹ in *K. rhizophila* and 28.395 µg mL□¹ in *G. arilaitensis* (Cipriani et al., unpublished data).

### Seed inoculation

Seedlings were transferred to fresh solid ½ MS plates at a density of five seeds per plate. Bacteria were grown in Plate Count liquid medium on the day before inoculation, and cell density was estimated spectrophotometrically at OD600 and adjusted to an optical density corresponding to approximately 10⁶ colony-forming units (CFU) mL□¹. Bacteria were pelleted by centrifugation (15,000 rpm, 5 min) and resuspended in ½ MS liquid medium to minimise osmotic stress upon transfer. Seedlings were then inoculated with 20 µL of the corresponding bacterial suspension in correspondence of the root; control seeds received medium only. Plates were sealed and incubated under the growth conditions described above for 7 days.

### Root phenotype analysis

Seedlings were photographed at 7 days post-inoculation (dpi). Images were analysed using ImageJ and the SmartRoot plugin for semi-automated root phenotyping [44,45]. Primary root length, root diameter, and the number of lateral roots were quantified on 15 seedlings per condition. Lateral root density was calculated as the number of lateral roots divided by primary root length and expressed as lateral roots cm□¹. Data visualisation and statistical analyses were performed in R (v4.4.1) and RStudio (v2024.12.0) (https://www.r-project.org/; https://www.posit.co/). For statistical analysis, the Kruskal–Wallis test followed by Dunn’s post hoc tests with Bonferroni correction were applied. Statistical significance was set at p < 0.05.

### RNA sequencing and sample preparation

For each standardized inoculum, five seedlings were pooled to have enough material and constituted one biological replicate, and three replicates per treatment were processed. Roots and shoots were separated with a razor bladeand flash-frozen in liquid nitrogen, homogenised with a Bullet Blender homogeniser. Total RNA was extracted using the CTAB method [46]. RNA yield and purity were assessed with a NanoDrop ND-1000 spectrophotometer (Thermo Fisher Scientific, MA, USA). RNA samples were sequenced by Novogene (UK) Co. The provider performed RNA quality control, poly(A)-enriched mRNA library preparation, sequencing on an Illumina NovaSeq X Plus platform (paired-end 150 bp; ∼12 Gb raw data per sample), and data quality control.

### Transcriptome analysis

Raw sequencing data were deposited in the NCBI SRA under BioProject PRJNA1472122 and processed on the Galaxy web platform (v25.0) using the public server at usegalaxy.org [47]. Adapters and low-quality bases were removed with Trimmomatic v0.39 [48] using the following parameters: ILLUMINACLIP:TruSeq3-PE.fa:2:30:10, LEADING:3, TRAILING:3, HEADCROP:10, SLIDINGWINDOW:4:15, and MINLEN:50. Reads were aligned to the tomato cv. Micro-Tom reference genome with RNA STAR v2.7.11b [49]. For samples sequenced across multiple lanes, BAM files were merged with samtools merge v1.18 [50]. Gene-level counts were obtained with featureCounts v2.1.1 [51]. Quality metrics were assessed with FastQC v0.74 and summarised with MultiQC v1.34 (https://www.bioinformatics.babraham.ac.uk/projects/fastqc/) [52]. Differential expression was analysed with DESeq2 v3.23 [53] in R (v4.4.1). Plots were generated using plotPCA, EnhancedVolcano v3.20, ggplot2 v4.0.3, and pheatmap v1.0.13. The Micro-Tom genome and annotation files were obtained from Xue et al. [54]. DEGs were filtered using thresholds of adjusted p-value ≤ 0.05 followed by |log2 FC| ≥ 1.

### Functional analysis

Functional annotation of differentially expressed genes (DEGs) was conducted against the KEGG (Kyoto Encyclopedia of Genes and Genomes) database. Analyses were performed in R (v4.4.1) using tidyverse v2.0.0, readxl v1.4.4, and KEGGREST v1.46.0. DEG identifiers from DESeq2 were cleaned to remove version suffixes and inner-joined to a gene-to-KEGG Orthology (KO) mapping table (https://eplant.njau.edu.cn/microTomBase/). KOs were linked to KEGG pathways with keggLink (KEGGREST). Global KEGG pathways were used as reference dataset. Non-plant pathways were excluded by keyword filtering. For each pathway, up- and down-regulated KOs were counted; a net difference metric (up- minus down-regulated KOs) was computed, and pathways exceeding 1.5 × SD of this metric were considered prominent in the dataset. Transcripts annotated as belonging to transcription factor (TF) families were isolated and analysed using NCBI BLAST to confirm the annotation and retrieve the best matching NCBI entry.

### Collection of plant tissues for metabolomic analysis

For metabolite extraction, five replicates consisting of seedlings were prepared 7 days after treatment. Roots and shoots were separated, flash-frozen in liquid nitrogen, and ground to a fine powder using a mortar and pestle. 100 mg of fresh tissue per sample was collected and lyophilised using an Edwards Freeze Dryer (Progitec MTP, Latina, Italy).

### Collection of plant exudates

At 7 dpi, 5 replicates consisting of 5 seedlings foreach treatment were transferred to 5 mL of 1 mM CaCl₂ to reduce osmotic stress on roots and kept in the growth chamber for 4 h to collect exudates. Exudate solutions were filtered through 0.22-µm polycarbonate membranes to remove bacterial cells, frozen at −80 °C, and lyophilised (Edwards Freeze Dryer; Progitec MTP, Latina, Italy).

### Collection of bacterial metabolites

For each al strain, five independent Plate Count liquid cultures were grown at 28 °C in the dark with shaking at 180 rpm. Upon reaching 10⁶ CFU mL□¹, cells were harvested by centrifugation (15,000 rpm, 5 min), resuspended in 2 mL ½ MS liquid medium, and incubated overnight. Culture supernatants were filtered through 0.22-µm polycarbonate filters to remove cells, frozen at −80 °C, and lyophilised with an Edwards Freeze Dryer (Progitec MTP, Latina, Italy).

### Metabolite extraction

Metabolite extraction and derivatisation [55] were carried out on lyophilised plant material, root exudates, and bacterial brothsby adding 1,400 µL of prechilled methanol (−20 °C) along with 60 µL of a 0.2 mg/mL ribitol solution in water, which served as an internal standard for the polar fraction. Tubes were vortexed for 10 s, then incubated at 70 °C with shaking (950 rpm) for 10 min in a thermomixer. Samples were centrifuged at 11,000 g for 10 min, and the resulting supernatant was transferred to glass vials. To each vial, 750 µL of cold chloroform (−20 °C) and 1,500 µL of ultrapure water (4 °C) were added sequentially, vortexed for 10 s, and then centrifuged at 2,200 g for 15 min. A 150 µL aliquot of the upper (polar) phase was collected, transferred to a 1.5 mL tube, and dried under vacuum without heat. Next, 40 µL of methoxyamine hydrochloride in pyridine (20 mg/mL) was added, and samples were incubated for 2 h at 30 °C (950 rpm). Silylation was performed by adding 70 µL of MSTFA and shaking for 30 min at 37 °C. The final derivatised volume (110 µL) was transferred into autosampler vials for GC–MS analysis.

### GC–MS conditions, analysis and data acquisition

One microlitre of each derivatised extract was injected in splitless mode onto a MEGA-5MS column (30 m × 0.25 mm × 0.25 µm + 10 m of precolumn) using a gas chromatograph (Agilent 7890A GC, Agilent Scientific Instruments, Santa Clara, CA, USA) coupled with a single quadrupole mass spectrometer (Agilent 5975C, Agilent Scientific Instruments, Santa Clara, CA, USA). Injector and ion source temperatures were maintained at 250 °C and 260 °C, respectively. The carrier gas (helium) flow was set to 1 mL/min. The oven program began with a 5 min hold at 70 °C, ramped at 5 °C/min to 350 °C, and ended with a 5 min hold at 330 °C. Mass spectra were acquired in electron-impact mode (70 eV) over an m/z range of 40–600 with a scan time of 0.2 s; the solvent delay was 9 min. Quality control injections, including pooled samples, n-alkane standards, and pyridine blanks, were interspersed throughout the sequence to monitor instrument performance, aid compound identification, and check retention index stability. We processed our data using MS-DIAL version 5.5 along with its open-source EI spectral library. The workflow consisted of raw peak extraction, baseline subtraction and calibration, chromatogram alignment, deconvolution, compound annotation, and peak-height integration [56]. Peak detection employed an average width of 20 scans and a minimum intensity threshold of 1,000 counts. During deconvolution, we applied a sigma window of 0.5 and discarded EI spectra with fewer than 5,000 counts. For compound identification, we allowed a retention-time window of ±0.2 min, an m/z tolerance of ±0.5 Da, required at least 60% EI spectral similarity, and imposed an overall identification score threshold of 80%. Alignment parameters were set with a 0.5 min retention-time tolerance and a tolerance factor of 0.5. Annotations were confirmed by matching against public libraries, including the Golm Metabolome Database’s MSRI spectra [57], MassBank [58], and MassBank of North America (MoNA, https://mona.fiehnlab.ucdavis.edu/), by comparing our EI patterns to those references. Metabolite assignments adhered to MSI guidelines [59], with Level 2 identifications based on database matches exceeding an 80% match factor.

### Metabolite analysis

Statistical analyses were conducted with the web-based platform MetaboAnalyst v6.0 (https://dev.metaboanalyst.ca/ModuleView.xhtml). For each condition, treated samples were compared against the corresponding control. Raw intensities were normalised to the internal standard (ribitol), log10-transformed, and Pareto-scaled. Univariate differences in metabolite abundance were assessed using two-sided t-tests with a significance threshold of p ≤ 0.05. Multivariate analyses comprised unsupervised principal component analysis (PCA) and supervised orthogonal projections to latent structures–discriminant analysis (OPLS-DA), both run with default parameters. To avoid overfitting, the OPLS-DA model was validated using permutation testing (n = 20 permutations). The model was considered robust when the permuted models showed significantly lower R² and Q² values than the original model, with empirical p-values ≤ 0.05. Additionally, the Q² intercept was negative, and the difference between R² and Q² values in the original model was within acceptable limits, indicating good predictive ability without overfitting. Discriminant metabolites contributing to class separation were identified using variable importance in projection (VIP) scores, with features showing VIP > 1 retained as discriminant features. Heatmaps were generated to visualise global metabolic patterns across treatments. Pathway analysis was also performed in MetaboAnalyst v6.0. Metabolite names were verified and curated against PubChem and the NIST Chemistry WebBook, then mapped to KEGG identifiers. The same normalisation pipeline (ribitol normalisation, log_10_ transformation, Pareto scaling) was applied prior to pathway enrichment. Because *Solanum lycopersicum* was not available among the listed species, *Arabidopsis thaliana* was used as the reference organism. Pathways were filtered at a false discovery rate (FDR) < 0.05 and ranked by pathway impact (range 0 to 1, with 1 indicating highest impact). Similarly, untargeted metabolomic profiles were acquired for all the culture broth samples. Raw metabolite peak intensities were normalised to ribitol as an internal standard to yield relative abundance values. These data were baseline-processed to generate a metabolite intensity matrix, which was subsequently normalised, log-transformed, and auto-scaled (mean-centred and divided by the standard deviation of each variable) prior to multivariate analysis. Univariate analysis was performed using one-way analysis of variance (ANOVA) across all detected features, reporting the F-statistic, raw parametric p-values, and false discovery rate (FDR)–adjusted p-values (Benjamini–Hochberg correction). For features that reached statistical significance, post hoc pairwise comparisons between conditions were conducted using Fisher’s least significant difference (LSD) test. Multivariate analysis comprised both unsupervised and supervised approaches. Unsupervised principal component analysis (PCA) was applied to assess global sample clustering, generating sample scores (PC1–PC15) and the corresponding metabolite loadings. Group separation in the PCA space was further evaluated using pairwise permutational multivariate analysis of variance (PERMANOVA) on the PCA scores, reporting pseudo-F statistics (F.Model), raw p-values, and adjusted p-values (p.adj). Supervised partial least squares–discriminant analysis (PLS-DA) was then performed to model class discrimination, yielding sample scores along latent variables (components 1–8), metabolite loading weights, variable importance in projection (VIP) scores, and regression coefficients (coef.mean).

## Data availability

The sequence dataset generated during the current study is available in the NCBI SRA database under BioProject PRJNA1472122. All datasets generated by the transcriptomic and metabolomic analyses are available as supplementary material on Zenodo (DOI: https://doi.org/10.5281/zenodo.20838081).

## Acknowledgements

We thank Nicole Zerbetto, MSc for her contribution during sampling, Dr. Davide Bulgarelli, University of Dundee (UK), Professor Yasin Dagdas, University of Heidelberg (DE), doctoral advisors of GG, for their valuable suggestions. We also thank Dr. Eunsook Park, University of Wyoming (USA), for helpful discussions, and Dr. Cecilia Zumajo Cardona, NYBG (USA), for sharing resources from the University of Padova STARS project SeedDive.

This work was supported by the Department of Biology PRID programme [grant BIRD225241 to SN]; the European Union’s Horizon 2020 research and innovation programme under the Marie Sklodowska-Curie Grant Agreement no 101034319 and from the European Union – NextGenerationEU [PhD fellowship grant to GG]; the Uni-Impresa 2020 programme of the University of Padova [Rep. 3350/2022 to BB]; and the Italian Ministry of University and Research under the European Union – NextGenerationEU initiative [PNRR Missione 4 Componente 2, “Dalla ricerca all’impresa”, Investimento 1.4, Progetto CN00000033 to BB]. RC is the recipient of a PhD fellowship DM117/2023 from the University of Padova and S.E.S.A. S.p.A., Este (Italy).

## Author contributions

Conceptualization, S.N. and G.G.; methodology, G.G., F.A. and S.N.; formal analysis, G.G. and F.A.; investigation, G.G., R.C. and F.A.; resources, S.N., B.B., F.A., T.B. and W.Z.; data curation, G.G. and F.A.; supervision, S.N. and B.B.; funding acquisition, S.N. and B.B.; writing—original draft preparation, G.G.; writing—review and editing, S.N., R.C., B.B. and F.A. All authors reviewed and approved the final version of the manuscript.

## Additional information

Competing interests: The authors declare no conflict of interest. Scientists from S.E.S.A. S.p.A. company, which provided material and funding, played no role in the study’s design, data collection, analysis, or interpretation, nor in the writing of the manuscript or the decision to publish the results.

## Supplementary figure legends

Figure S1. Bacterial inoculation differentially affects tomato root architecture at 7 days post-inoculation. (a–c) Primary root length, lateral root number and lateral root density in non-inoculated control seedlings and seedlings inoculated with *Bacillus sonorensis*, *Glutamicibacter arilaitensis* or *Kocuria rhizophila*. Primary root length is expressed in cm, lateral root number represents the absolute number of lateral roots per seedling, and lateral root density is expressed as the number of lateral roots per cm of primary root. (d) Primary root diameter in control and inoculated seedlings. n = 15 seedlings per condition. Statistical significance was assessed using the Kruskal–Wallis test followed by Dunn’s post hoc test with Bonferroni correction. Asterisks indicate significant differences from the control: P < 0.05, **P < 0.001 and ***P < 0.0001.

Figure S2. Global transcriptomic profiles differ among bacterial treatments in tomato shoots and roots. Principal component analysis (PCA) of shoot (a) and root (b) transcriptomes, including control samples and samples inoculated with *Bacillus sonorensis*, *Kocuria rhizophila* or *Glutamicibacter arilaitensis* in the same analysis. n = 3 biological replicates per condition; each replicate comprised shoot or root tissue pooled from five seedlings.

Figure S3. Bacterial treatments induce distinct gene-expression patterns in tomato shoots and roots. Heatmaps of differentially expressed genes (DEGs) in shoots (a–c) and roots (d–f) at 7 days post-inoculation with *Bacillus sonorensis*, *Kocuria rhizophila* and *Glutamicibacter arilaitensis*, respectively, relative to non-inoculated controls. DEGs were selected using thresholds of adjusted P value ≤ 0.05 and |log2 fold change| ≥ 1. Heatmap colours indicate relative gene-expression values. n = 3 biological replicates per condition; each replicate comprised shoot or root tissue pooled from five seedlings.

Figure S4. Bacterial inoculation produces isolate- and tissue-specific changes in primary-metabolite abundance. Heatmaps of significantly altered metabolites in shoots (a–c) and roots (d–f) of seedlings inoculated with *Bacillus sonorensis*, *Kocuria rhizophila* and *Glutamicibacter arilaitensis*, respectively, at 7 days post-inoculation. Only metabolites differing significantly from the corresponding non-inoculated controls are shown. Heatmap colours indicate relative metabolite abundance after data normalisation and scaling. n = 5 biological replicates per condition; each replicate comprised shoot or root tissue pooled from five seedlings. Analyses were performed using MetaboAnalyst 6.0.

Figure S5. Multivariate analyses reveal distinct metabolomic profiles in tomato shoots and roots following bacterial inoculation. Principal component analysis (PCA) and orthogonal partial least squares–discriminant analysis (OPLS-DA) of shoot (a,b) and root (c,d) samples, including controls and the three bacterial treatments. n = 5 biological replicates per condition; each replicate comprised shoot or root tissue pooled from five seedlings. Analyses were performed using MetaboAnalyst 6.0.

Figure S6. Relative abundances of metabolites contributing to treatment separation in tomato shoots. Metabolites were selected using a variable importance in projection (VIP) score > 1. Panel a shows metabolites displaying a common response across the three bacterial treatments relative to the control. Panels b–d show treatment-specific metabolites for *Bacillus sonorensis*, *Kocuria rhizophila* and *Glutamicibacter arilaitensis*, respectively. n = 5 biological replicates per condition; each replicate comprised shoot tissue pooled from five seedlings.

Figure S7. Relative abundances of metabolites contributing to treatment separation in tomato roots. Metabolites were selected using a variable importance in projection (VIP) score > 1. Panel a shows metabolites displaying a common response across the three bacterial treatments relative to the control. Panels b–d show treatment-specific metabolites for *Bacillus sonorensis*, *Kocuria rhizophila* and *Glutamicibacter arilaitensis*, respectively. n = 5 biological replicates per condition; each replicate comprised root tissue pooled from five seedlings.

Figure S8. Bacterial inoculation induces isolate-specific changes in tomato root-exudate composition. Heatmaps of significantly altered metabolites in root exudates collected from seedlings inoculated with *Bacillus sonorensis* (a), *Kocuria rhizophila* (b) or *Glutamicibacter arilaitensis* (c) at 7 days post-inoculation. Only metabolites differing significantly from the corresponding non-inoculated controls are shown. Heatmap colours indicate relative metabolite abundance after data normalisation and scaling. n = 5 biological replicates per condition; each replicate comprised root exudates collected from five seedlings. Analyses were performed using MetaboAnalyst 6.0.

Figure S9. Multivariate analyses distinguish root-exudate profiles associated with the three bacterial treatments. Principal component analysis (PCA; a) and orthogonal partial least squares–discriminant analysis (OPLS-DA; b) of root exudates from control seedlings and seedlings inoculated with *Bacillus sonorensis*, *Kocuria rhizophila* or *Glutamicibacter arilaitensis*. n = 5 biological replicates per condition; each replicate comprised root exudates collected from five seedlings. Analyses were performed using MetaboAnalyst 6.0.

Figure S10. Relative abundances of root-exudate metabolites contributing to treatment separation. Metabolites were selected using a variable importance in projection (VIP) score > 1. Panel a shows metabolites displaying a common response across the three bacterial treatments relative to the control. Panels b–d show treatment-specific metabolites for *Bacillus sonorensis*, *Kocuria rhizophila* and *Glutamicibacter arilaitensis*, respectively. n = 5 biological replicates per condition; each replicate comprised root exudates collected from five seedlings.

Figure S11. Metabolite profiles of cell-free culture filtrates of the three bacterial isolates. (A) Principal component analysis (PCA) scores plot of culture filtrates from *Bacillus sonorensis*, *Kocuria rhizophila,* and *Glutamicibacter arilaitensis*. PC1 and PC2 explain 79.8% and 15.7% of the total variance, respectively. Shaded ellipses indicate 95% confidence regions. (B) PCA loadings plot showing the contribution of each metabolite to sample separation. (C) Hierarchical clustering heatmap of metabolites differing significantly among isolates by one-way ANOVA (FDR-adjusted p < 0.05). Each column shows the mean of one isolate. Colours indicate relative abundance after auto-scaling, from lower (blue) to higher (red). Peak intensities were normalised to ribitol, log10-transformed and auto-scaled. n = 5 independent cultures per isolate. Complete results are provided in Supplementary File 1.

## Supplementary information

The following supplementary data are available at Zenodo (DOI https://doi.org/10.5281/zenodo.20838082)

Supplementary Tables S1-S14: SHOOT and ROOT UpSet gene lists with KEGG annotation

Supplementary Tables S15-S17 SHOOT Heatmaps with KEGG annotation for each inoculum

Supplementary Tables S18-20 ROOT Heatmaps with KEGG annotation for each inoculum

Supplementary Table S21. DEG TF genes in the different treatments

Supplementary Tables S22-24 Metabolomic of Shoot for each inoculum.xlsx

Supplementary Tables S25-27 Metabolomic of Root for each inoculum.xlsx

Supplementary Tables S28-30 Metabolic pathways analysis of Shoot for each inoculum.xlsx

Supplementary Tables S31-33 Metabolic pathways analysis of Root for each inoculum.xlsx

Supplementary Tables S34-36 Metabolomic analysis of root exudates for each inoculum.xlsx

Supplementary File 1 Metabolomic analysis of bacterial culture broths

