## Supplementary figures and images for "Transcriptomic and metabolomic profiling reveals early responses of tomato plants elicited by compost-derived bacteria"

### Supplementary Fig. S1

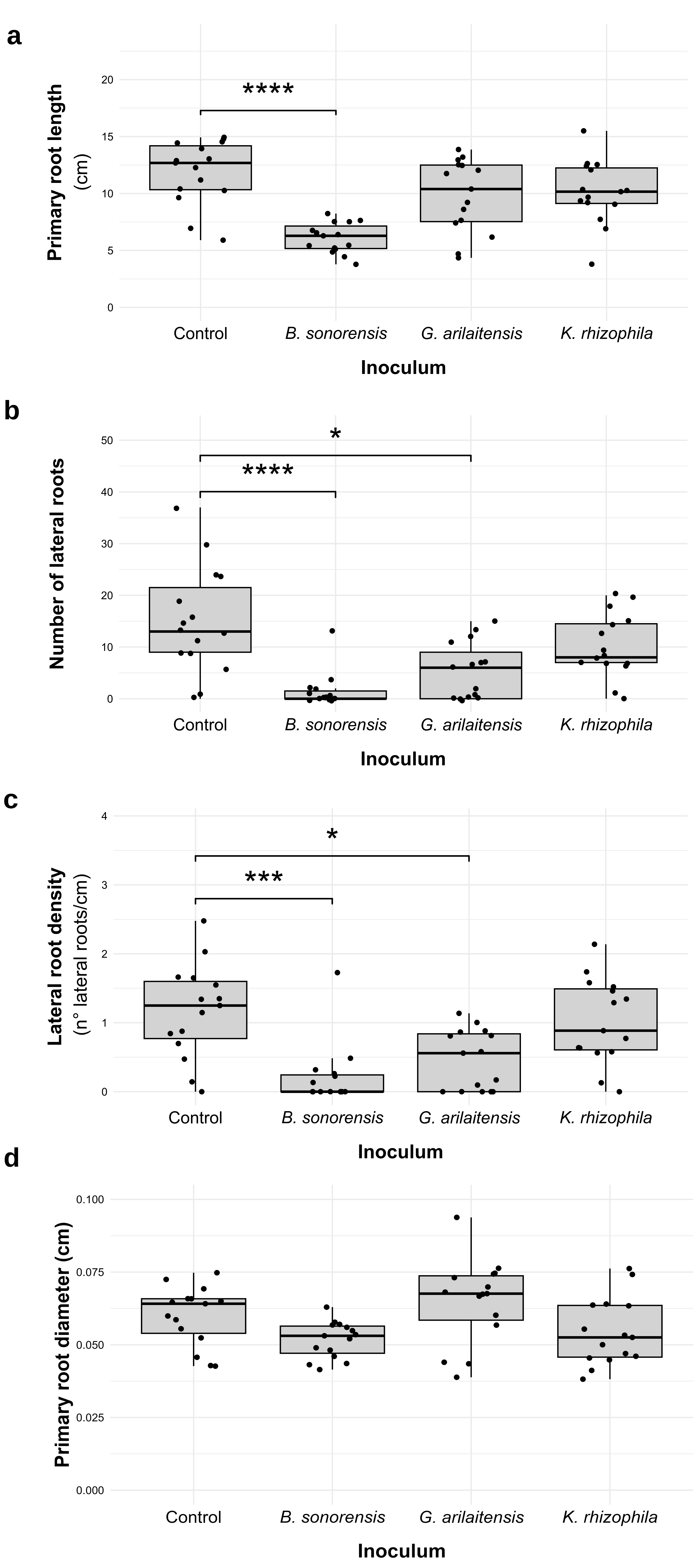

### Supplementary Fig. S2

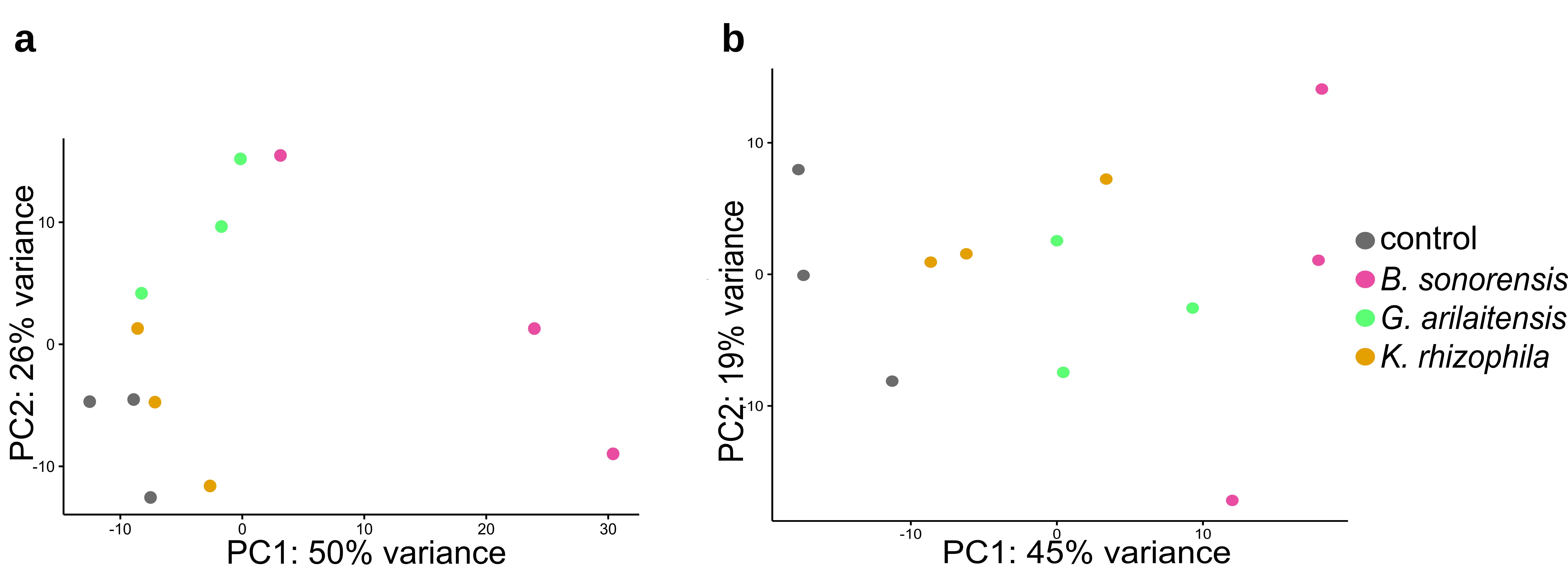

### Supplementary Fig. S3

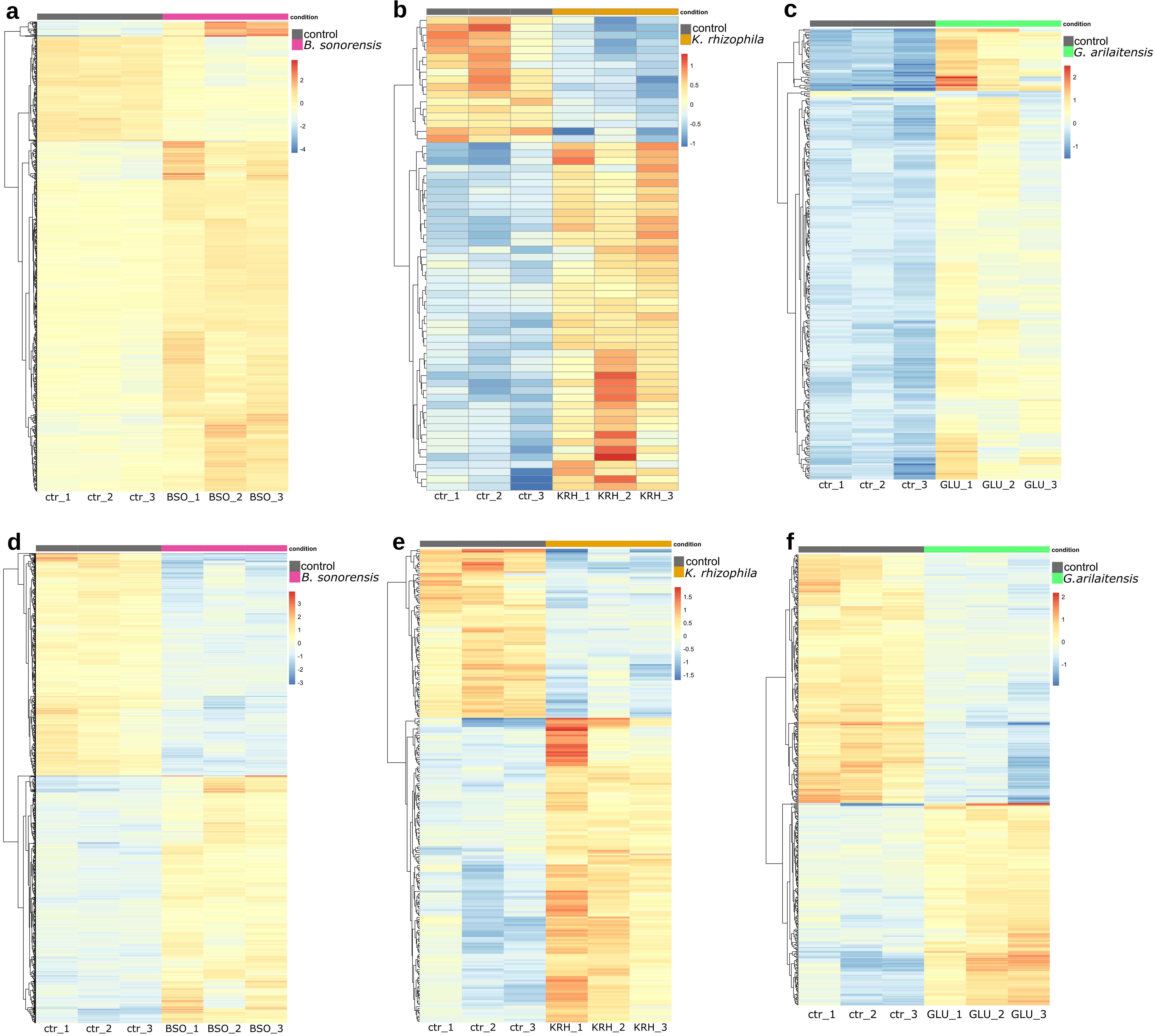

### Supplementary Fig. S4

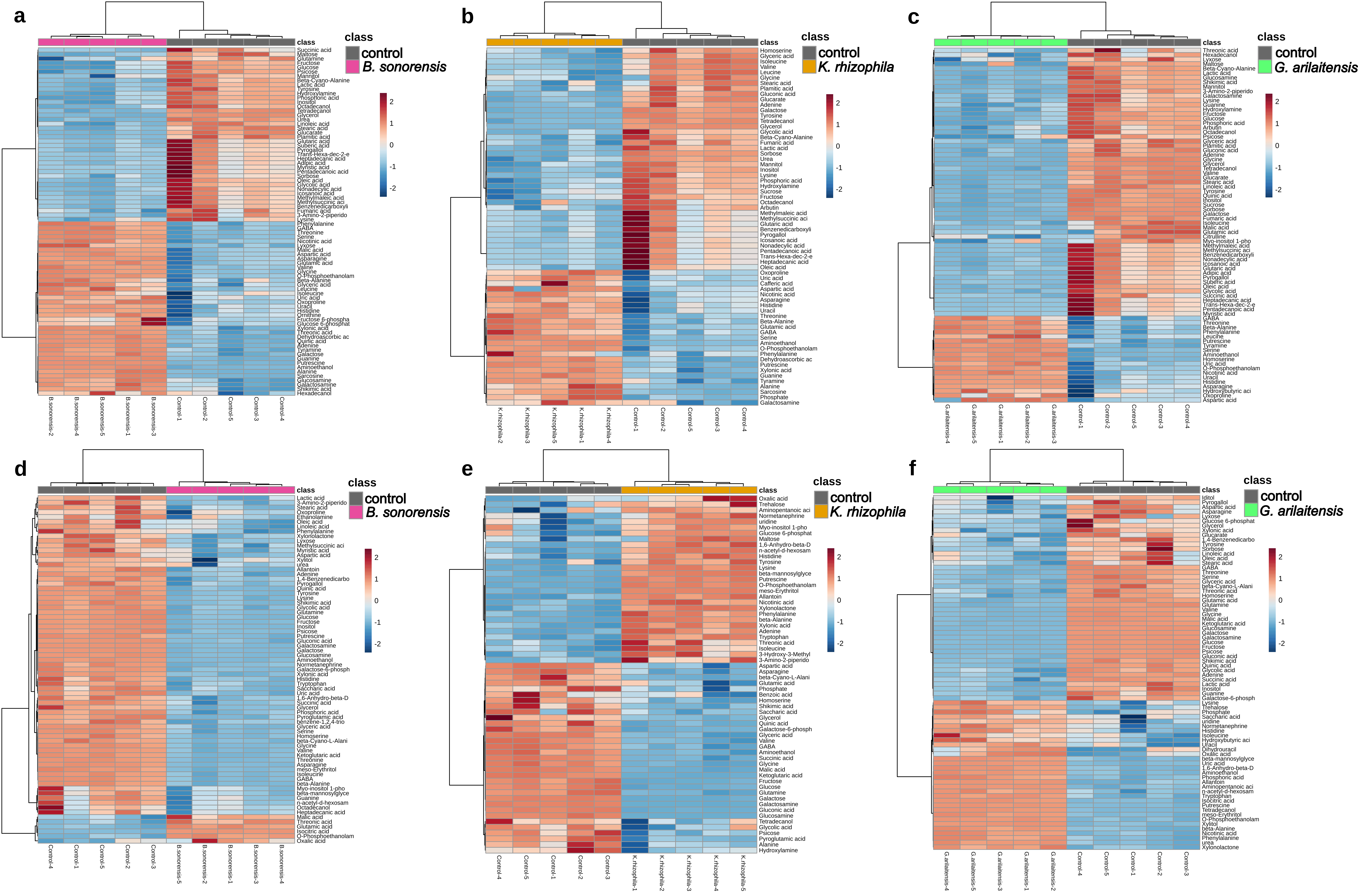

### Supplementary Fig. S5

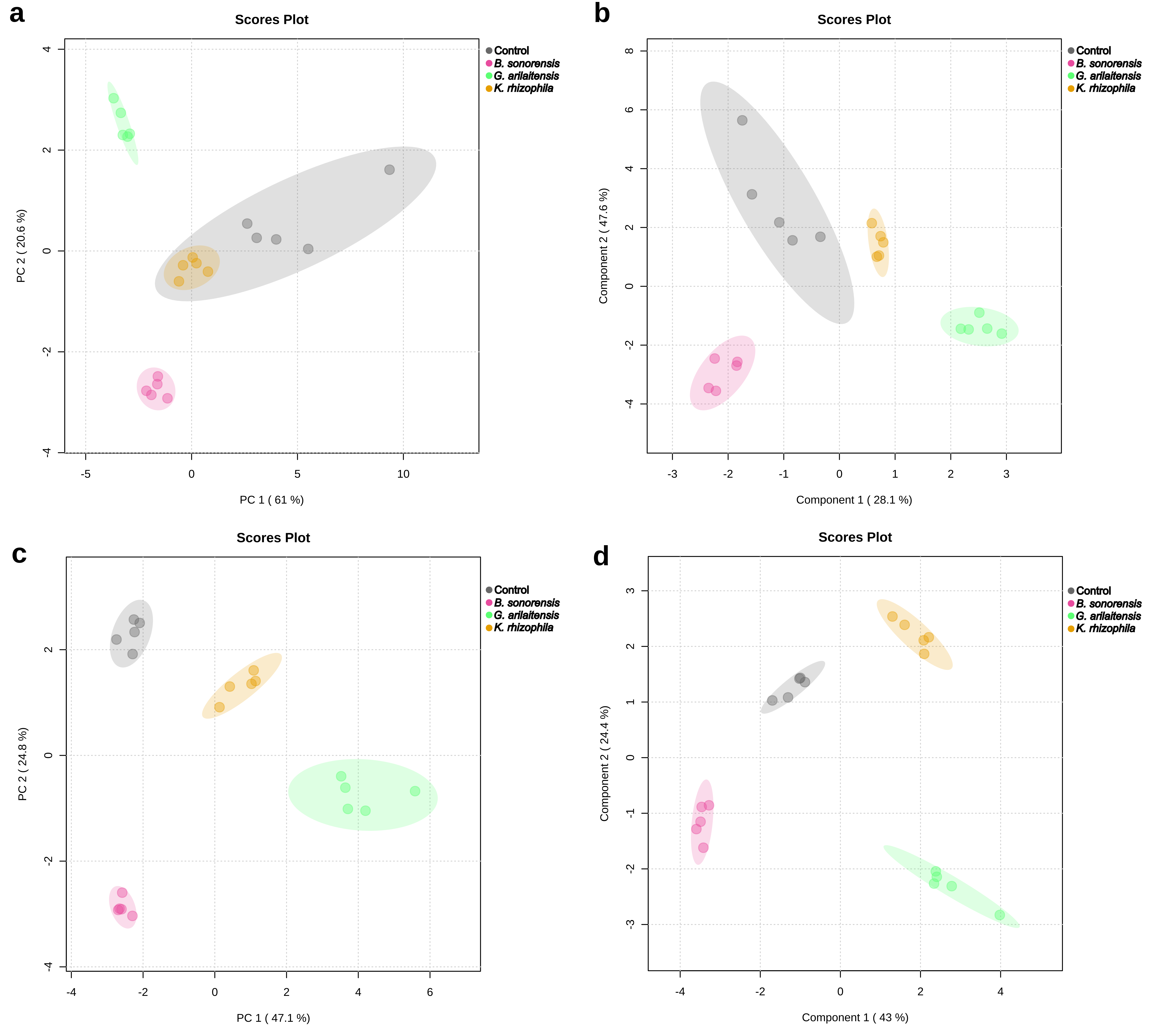

### Supplementary Fig. S6

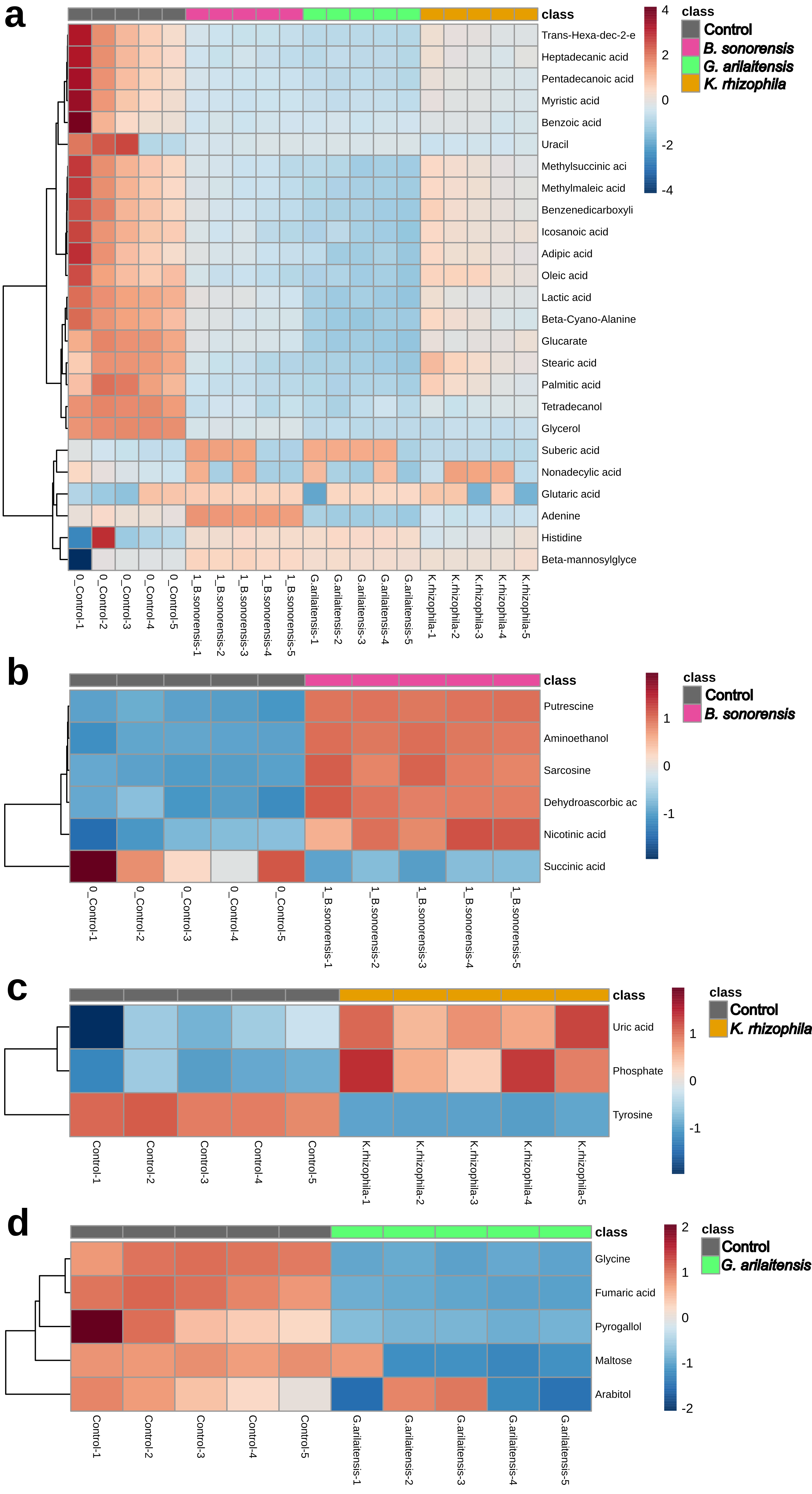

### Supplementary Fig. S7

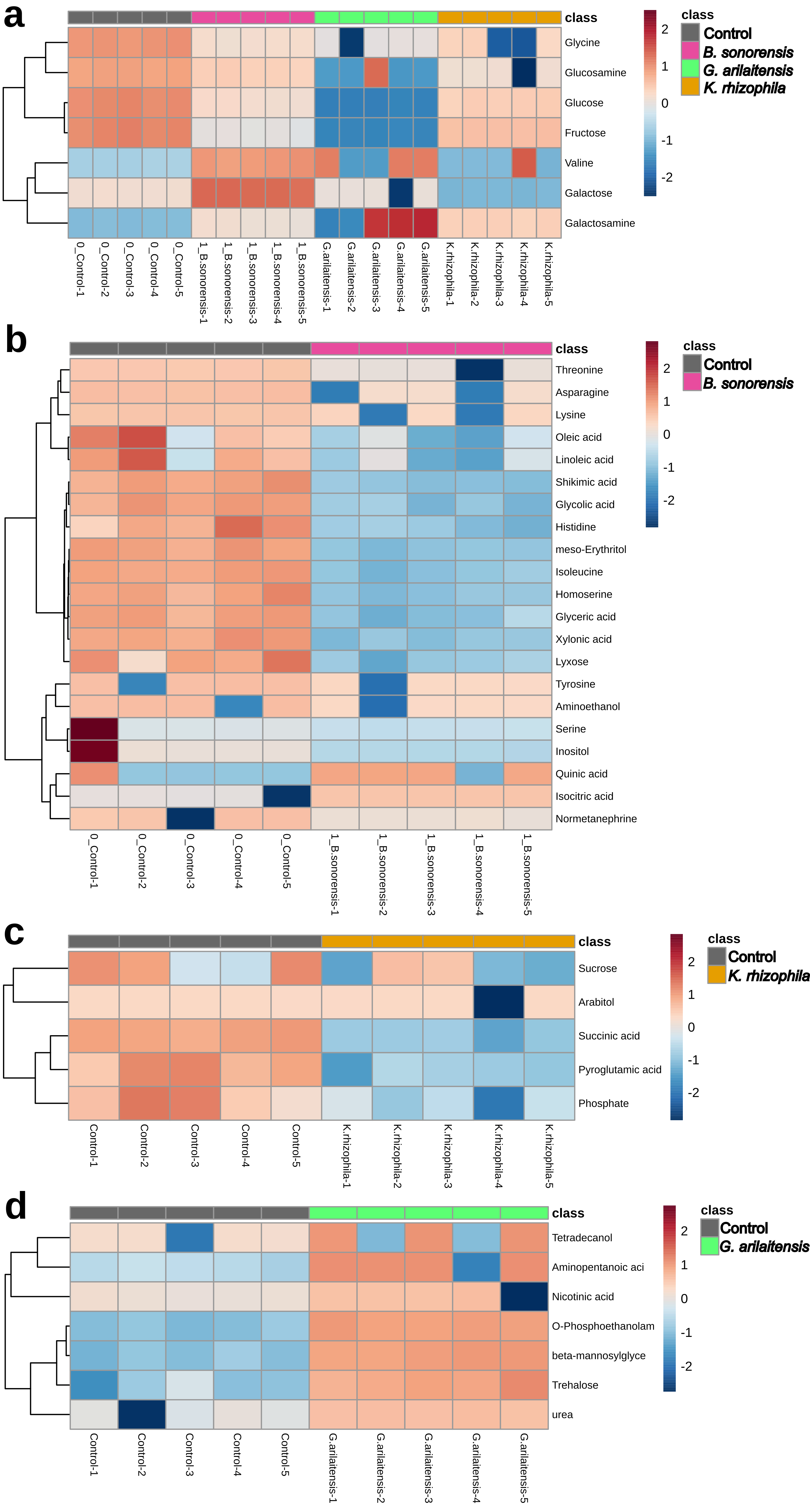

### Supplementary Fig. S8

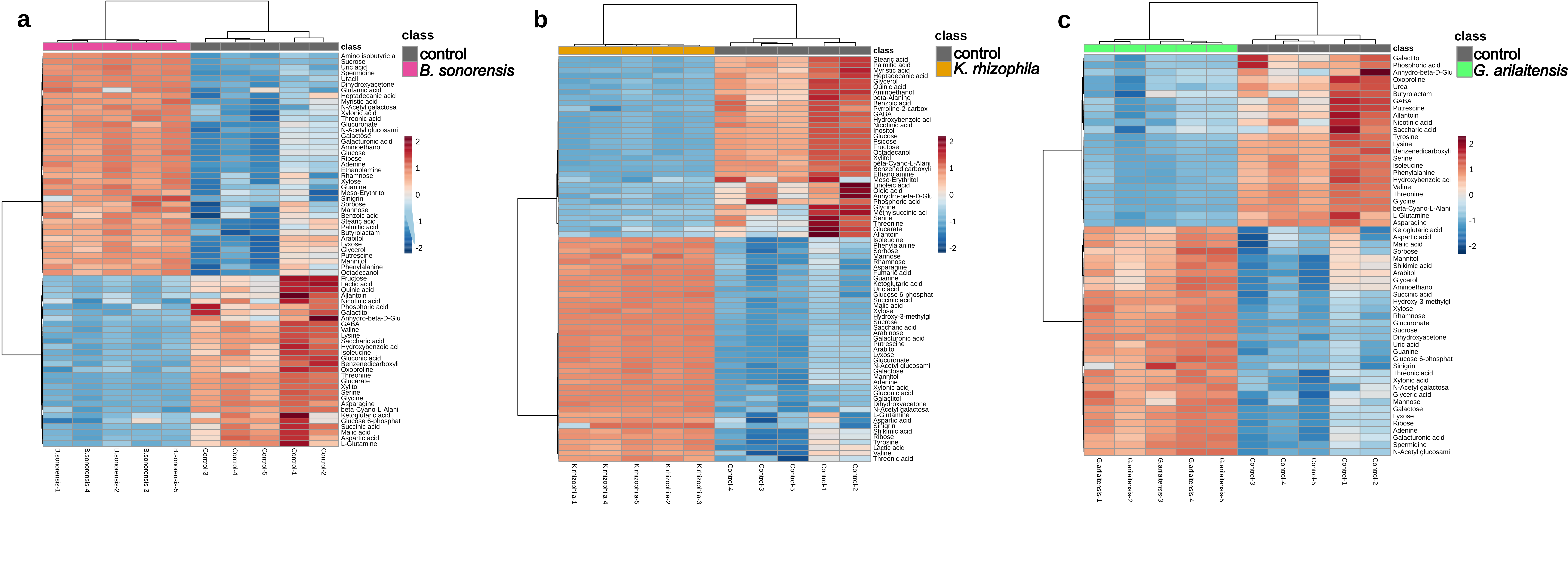

### Supplementary Fig. S9

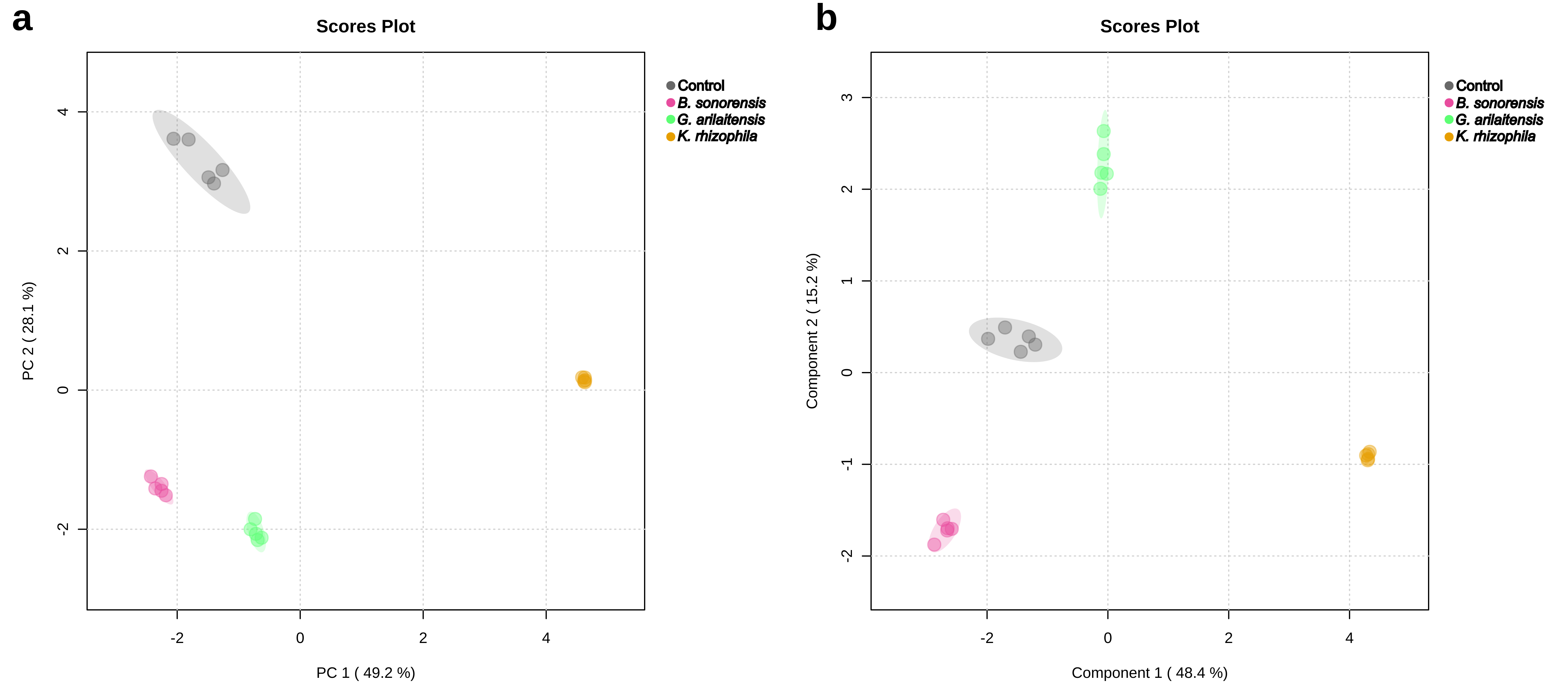

### Supplementary Fig. S10

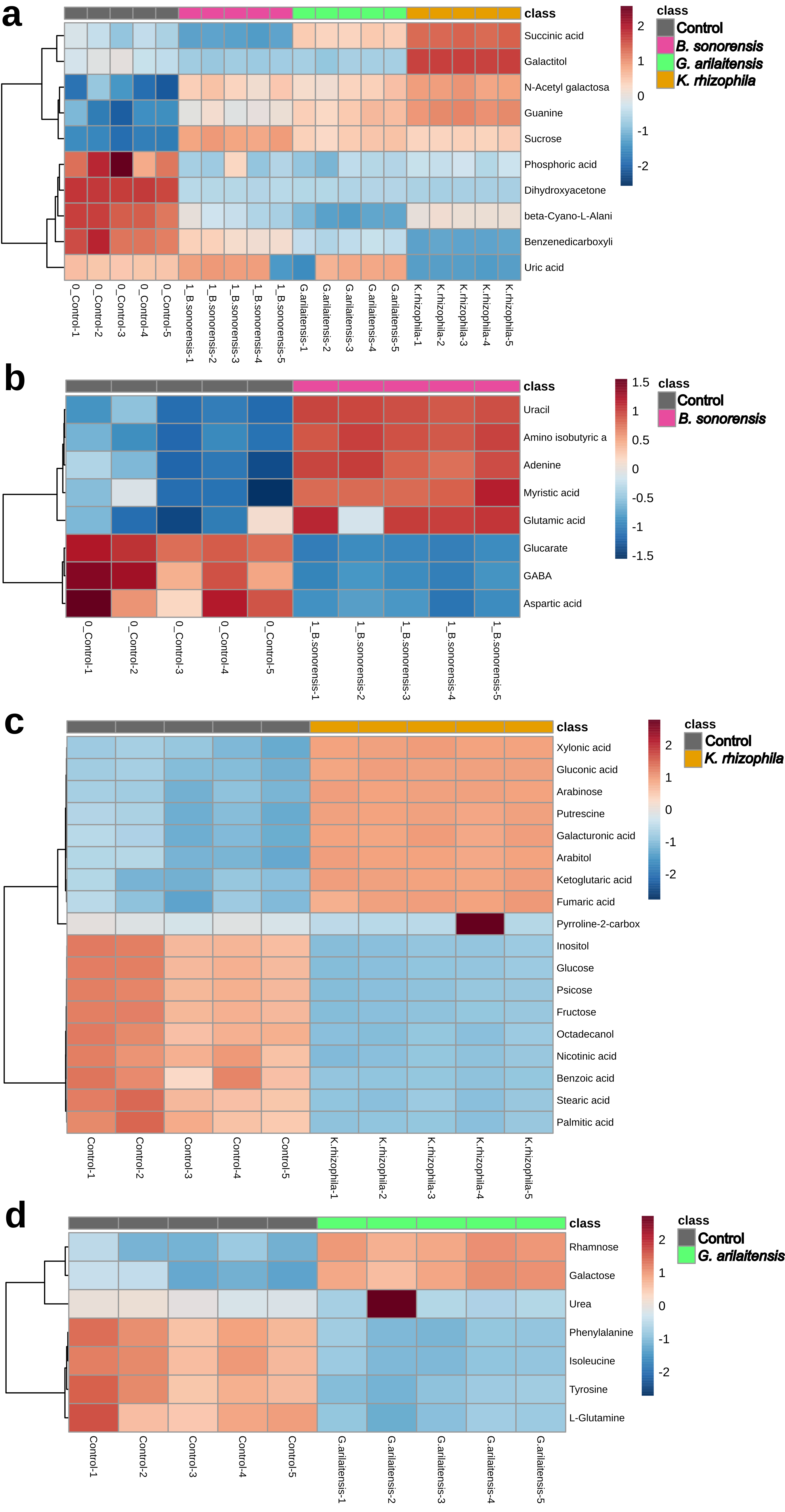

### Supplementary Fig. S11

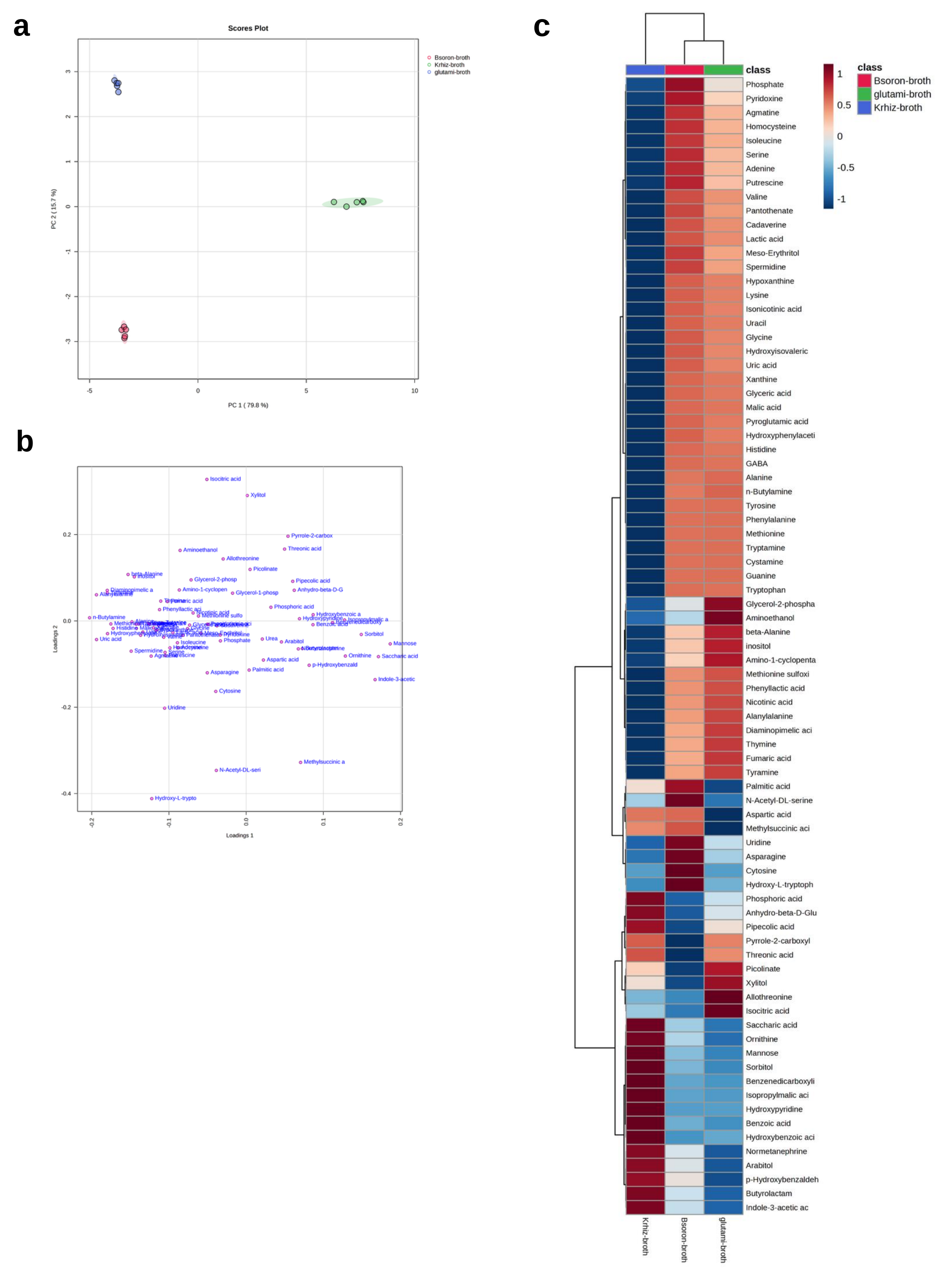
